# Multi-omic mapping of *Drosophila* protein secretomes reveals tissue-specific origins and inter-organ trafficking

**DOI:** 10.1101/2025.07.09.659702

**Authors:** Justin A. Bosch, Pierre Michel Jean Beltran, Cooper Cavers, James Thai LaGraff, Randy Melanson, Ankita Singh, Weihang Chen, Yanhui Hu, Sudhir Gopal Tattikota, Ying Liu, Yousuf Hashmi, John M. Asara, Tess Branon, Alice Y. Ting, Steven A. Carr, Norbert Perrimon

**Affiliations:** Department of Genetics, Blavatnik Institute, Harvard Medical School, Boston, MA, USA; Department of Human Genetics, University of Utah School of Medicine, Salt Lake City, UT, USA; Broad Institute of MIT and Harvard, Cambridge, MA, USA; Division of Signal Transduction, Beth Israel Deaconess Medical Center, Boston, MA, USA; Department of Medicine, Harvard Medical School, Boston, MA, USA; Departments of Genetics, Biology, and by courtesy, Chemistry, Stanford University, Stanford, CA, USA; Department of Chemistry, Massachusetts Institute of Technology, Cambridge, MA, USA; Chan Zuckerberg Biohub – San Francisco, San Francisco, CA, USA; Howard Hughes Medical Institute, Boston, MA, USA

**Keywords:** Proximity labeling, secretome, *Drosophila*, TurboID, proteomics, blood proteins, snRNA-seq, CRISPR, inter-organ communication

## Abstract

Secreted proteins regulate many aspects of animal biology and are attractive targets for biomarkers and therapeutics. However, comprehensively identifying the “secretome”, along with their tissues of origin, remains extremely challenging. To address this, we employed multiple ‘omics methods to define a tissue-secretome map of 535 blood plasma proteins derived from specific cell-types and organs in *Drosophila melanogaster*. This map was enabled by methodological improvements including a collection of transgenic flies to label endogenous secreted proteins in 10 major tissue types, large-scale blood isolation, whole animal snRNA-seq, and a collection of 40 knock-in strains. Using this map, we discover surprising findings about circulating proteins: most originate from specific tissues including unusual sources (e.g. glia), many are uncharacterized, and some are shed ectodomains of transmembrane proteins. In addition, *in vivo* experiments revealed circulating proteins with remarkably tissue-specific expression, as well as proteins that are deposited in a different tissue from where they are synthesized, suggesting potential inter-organ functions. Our secretome map will serve as a resource to investigate blood protein function, discover novel tissue-tissue communication signals, and mine for homologues of human biomarkers.

## Introduction

Secreted proteins play essential roles in development, immunity, and metabolism by acting in the extracellular space to influence nearby or distant cells. Additionally, disruptions in secreted protein signaling are implicated in a wide range of diseases, including diabetes and cancer. Because they are more accessible than intracellular proteins, secreted proteins and their potential cell surface receptors are attractive targets for both therapeutics and biomarkers. While efforts to define the full set of secreted proteins (the “secretome”) have yielded thousands of candidates^1,2^, we still lack a comprehensive map of their cell-type-specific sources, extracellular distribution (local versus circulating), and non-autonomous functions. Such a detailed map of secreted proteins would enhance our understanding of basic biological processes and disease mechanisms, and lead to improved therapeutics and biomarkers.

Mapping secreted proteins remains technically challenging. For example, mass spectrometry (MS) of extracellular fluids can identify secreted proteins but not their source cells^3^. While the extraction of extracellular fluids from specific tissues can be performed before MS^4^, it is not adaptable to all cell-types. Furthermore, combining MS with RNA-seq has been used to map candidate tissue sources^2,5^, but this approach does not physically trace the secreted protein back to the origin tissue and cannot distinguish constitutive secretion from cell lysis or regulated secretion. Additional challenges of MS are that secreted proteins are obscured by abundant proteins or misattributed to the target tissues they accumulate in rather than those that produce them. These issues highlight the need for new tools, especially those that directly trace and enrich secreted proteins from complex biological samples.

Proximity labeling (PL) enzymes have recently emerged as a promising solution. Promiscuous biotin ligases like BioID and TurboID can label endogenous proteins in living cells with biotin^6,7^, and biotinylated proteins can be isolated using streptavidin beads. Thus, biotinylated proteins can be enriched from specific cell populations in complex biological samples, reducing background. When targeted to the secretory pathway, these enzymes can label endogenous secreted proteins, which are then recovered from the extracellular space. Several studies have used promiscuous biotin ligases to identify cell/tissue-specific secretomes^8–16^ as well as proteins trafficking between cells and organs^17,18^. However, these studies are limited to a few cell types or organs, and broader efforts are needed to approach a full tissue-level secretome map. Furthermore, while many of these studies utilize mouse models, systematic *in vivo* characterization of candidate secreted proteins remains challenging due to the time, cost, and complexity of functional studies in mammals.

*Drosophila* is well suited for this task due to its conserved physiology and powerful genetic tools. For example, flies have conserved organ systems with mammals, including brain, muscle, and gut. Furthermore, its blood (or hemolymph) is analogous to human plasma, containing human-conserved circulating proteins that control immunity, coagulation, and “inter-organ” signaling (e.g. hormones)^19–21^. *Drosophila* genetic tools also allow rapid *in vivo* characterization of candidate genes at large-scale, facilitated by an enormous collection of tissue-specific Gal4 driver transgenes^22–24^. Importantly, tissue-specific expression of proximity labeling enzymes in *Drosophila* are non-toxic^6,18,25,26^. However, its small volume of blood (or hemolymph) has prevented a large-scale unbiased secretome analysis. Therefore, improved protocols and tools are required.

Here, we employed TurboID proximity labeling, mass spectrometry, single nuclear RNA-seq (snRNA-seq), and CRISPR gene editing to create a comprehensive secretome map in *Drosophila* third instar larvae. We identified 535 circulating secreted proteins, and their tissues of origin, to discover new insights into the relative contributions of different tissues to blood proteins, identify uncharacterized tissue-specific secreted proteins, and reveal proteins that are synthesized in one tissue and localized to another, suggesting inter-organ functions. This work establishes a scalable platform for secretome discovery in *Drosophila* and opens the door to systematic exploration of inter-tissue signaling.

## Results

### Characterization of ER-localized TurboID to label secreted proteins in cultured *Drosophila* S2R+ cells

To identify the tissue-of-origin of secreted proteins in *Drosophila*, we sought to use an *in vivo* proximity labeling approach. Like other studies, we wanted to genetically express a promiscuous biotin ligase that is targeted to the endoplasmic reticulum (ER) lumen to label endogenous secreted proteins^11–15,18^, which are then purified from the extracellular space (**Figure 1A**). We previously labeled proteins in the secretory pathway using Myc-tagged promiscuous biotin ligase BirA*G3 in the ER (*Myc-BirA*G3-ER*)^6,11,18^. BirA*G3 (G3 = Generation 3) is a precursor to the engineered promiscuous biotin ligase TurboID (which has three additional mutations compared to G3), where TurboID has higher biotinylation activity than BirA*G3^6^. Therefore, we constructed updated plasmids to express ER-localized TurboID (*GFP-TurboID-ER*) (**Figure 1A**). For comparison, we constructed plasmids to express a cytoplasmic TurboID (*GFP-TurboID*).

**Figure 1:**
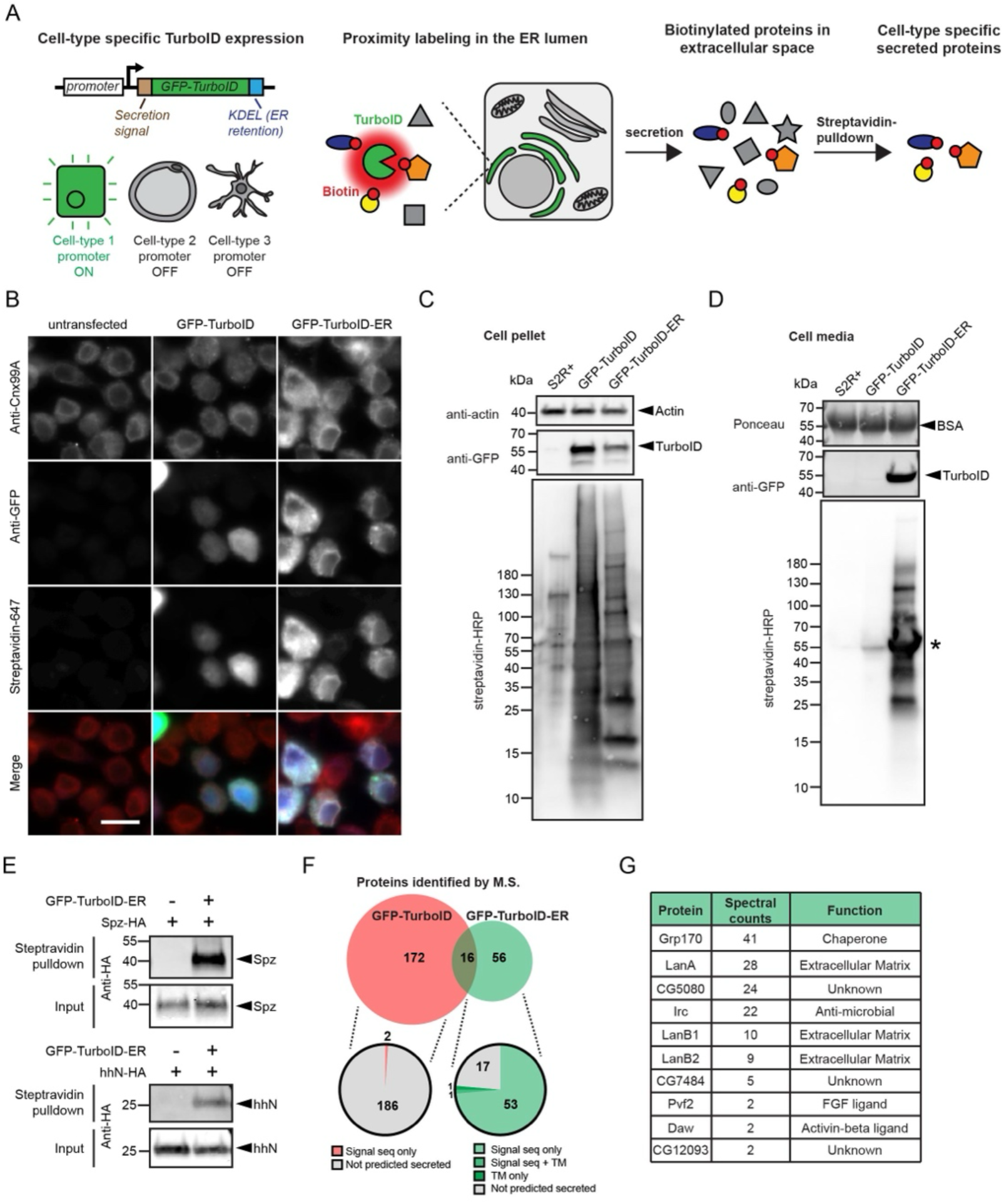
Labeling secreted proteins in cultured fly cells using ER-localized TurboID. **(A)** Schematic of genetic cell-type-specific labeling of endogenous proteins in the ER-lumen with GFP-TurboID, followed by purification of biotinylated proteins from the extracellular space. **(B)** Confocal microscopy of S2R+ cells transfected with *pMT-GFP-TurboID* or *pMT-GFP-TurboID-ER*. ER detected with Anti-Cnx99A (red), biotinylated proteins detected with streptavidin-647 (blue), and GFP-TurboID detected with anti-GFP (green). Scale bar is 10µm. Red and green channel signal intensity is the same for all samples. Blue channel signal for GFP-TurboID sample was lowered to match the signal intensity of GFP-TurboID-ER sample **(C-D)** Western blot of cell lysates **(C)** or cell media **(D)** from S2R+ cells stably expressing GFP-TurboID or GFP-TurboID-ER. Each lane loaded 15µg protein. Arrowheads indicate expected band, asterisk indicates auto-biotinylated GFP-TurboID. **(E)** Western blot of streptavidin-pulldown of HA-tagged secreted proteins in media from S2R+ cells transfected with *pMT-GFP-TurboID-ER*. Arrowheads indicate expected bands. Each lane loaded 15µg protein. Predicted molecular weight: Spz-HA 38.3 kDa, hhN-HA 23.6 kDa **(F)** Analysis of proteins identified following whole biotinylated protein pulldown from cell media, gel lane excision, and liquid chromatography-tandem mass spectrometry (LC-MS/MS). Venn diagram showing the number of proteins identified (top left), pie charts showing the fraction of predicted secreted proteins (bottom left) and table showing example secreted proteins identified from *pMT-GFP-TurboID-ER* cell media (right).

To rapidly test and characterize this TurboID labeling strategy, we first performed experiments in cultured *Drosophila* S2R+ cells. Notably, this TurboID labeling approach has not been applied to any cultured *Drosophila* cell line, so we saw an opportunity to establish a resource that could benefit the broader community, even though this was not our primary objective.

To test if GFP-TurboID-ER labels proteins in the secretory compartment, we transfected TurboID DNA constructs into S2R+ cells grown in biotin-supplemented media, and visualized GFP and biotinylated proteins with confocal microscopy. GFP-TurboID-ER fluorescent signal was excluded from cell nuclei, enriched in a punctate pattern surrounding the nucleus, and overlapped with the ER protein Calnexin 99A (**Figure 1B**). Biotinylated proteins were similarly localized to the ER (**Figure 1B**). In contrast, we detected GFP-TurboID and biotinylated proteins in the cytoplasm and nucleus (**Figure 1B**). We observed similar results using our previous Myc-BirA*G3 Myc-BirA*G3-ER constructs (**Supplemental Figure 1A**), indicating that our TurboID constructs are properly localized like prior BirA*G3 versions.

Next, we tested whether we could detect secreted biotinylated proteins in cell culture media supernatant from cells expressing GFP-TurboID-ER. To induce robust expression, we generated stable S2R+ cell lines expressing either GFP-TurboID or GFP-TurboID-ER under the control of a copper-inducible metallothionein (*MT*) promoter. After incubating cell lines with CuSO4 and excess biotin for 24 hours, we subjected cell pellets and cell media supernatant to SDS-PAGE, followed by western blotting using streptavidin-HRP to detect biotinylated proteins. For *MT-GFP-TurboID-ER* cells, we detected a smear of biotinylated protein bands of different molecular weights from both cell pellets and cell media (**Figures 1C-D**), suggesting that both intracellular and secreted proteins are being labeled. For *MT-GFP-TurboID* cells, we also saw a smear of biotinylated proteins from cell pellets, but with a different banding pattern compared to *MT-GFP-TurboID-ER* (**Figure 1C**). Unlike *MT-GFP-TurboID-ER* cells, biotinylated proteins in media from *MT-GFP-TurboID* cells were barely detectible, indicating minimal secretion of TurboID-labeled cytosolic proteins into the extracellular medium (**Figure 1D**). While we detected GFP-TurboID and GFP-TurboID-ER in cell pellets by anti-GFP westerns (**Figure 1C**), we also detected GFP-TurboID-ER in cell media supernatant (**Figure 1D**), suggesting that the protein can be secreted. Similarly, we observed a band corresponding to the same molecular weight as GFP-TurboID-ER (∼55 kDa) on streptavidin-HRP blots (**Figure 1D**), suggesting that TurboID auto-biotinylates, which has been previously described^6^. We observed similar results using transfection of *MT-Myc-BirA*G3-ER* in S2R+ cells (**Supplemental Figure 1B**).

Next, we tested if GFP-TurboID-ER biotinylates known secreted proteins and if we could recover them from culture media supernatant. To accomplish this, we transfected S2R+ cells with HA-tagged versions of two secreted ligands (Spatzle and Hedgehog), collected culture media, enriched biotinylated proteins on streptavidin beads, and detected the HA-tagged proteins by western blot. Whereas we observed both Spatzle-HA and Hedgehog-HA in input samples, they were only detected after pulldowns when co-transfecting *MT-GFP-TurboID-ER* (**Figure 1E**).

To further confirm that GFP-TurboID-ER biotinylates secreted proteins, we used liquid chromatography-tandem mass spectrometry (LC-MS/MS) to identify labeled proteins in culture media from stable cell lines. Using streptavidin beads to enrich biotinylated proteins in culture media supernatant, we ran the eluted proteins on an SDS-PAGE gel and detected total proteins using a Coomassie stain. We observed a smear of protein bands from *MT-GFP-TurboID-ER* cell media, which were not present from negative control wild-type S2R+ cell media, including a prominent band at the molecular weight of auto-biotinylated TurboID (**Supplemental Figure 1C**). Similar results were observed using BirA*G3 expressing cells **(Supplemental Figure 1D)**. Unexpectedly, we also observed a smear from *MT-GFP-TurboID* cell media pulldowns. This is despite gentle harvesting and filtering of cell media (see methods), suggesting that S2R+ cells undergo cell lysis during normal culture and/or unconventional secretion. To identify proteins labeled by GFP-TurboID or GFP-TurboID-ER, we cut out entire gel lanes, followed by in-gel digestion and LC-MS/MS. We identified 188 *Drosophila* proteins from *MT-GFP-TurboID* cell media and 72 from *MT-GFP-TurboID-ER* cell media (**Figure 1G, Supplemental Table 1A**). Proteins that are predicted as extracellular, such as containing a signal sequence and/or a TM domain, were highly enriched from GFP-TurboID-ER media (76%) relative to GFP-TurboID media (1%) (**Figure 1G, Supplemental Table 1A**). These secreted proteins include signaling proteins, extracellular matrix proteins, chaperones, and uncharacterized proteins (**Figure 1G**). We also identified GFP-TurboID in media from both cell lines, confirming the protein is released into the media (**Supplemental Table 1B**). Interestingly, we also identified cow proteins such as Complement C3, suggesting GFP-TurboID in media may label proteins in culture media derived from fetal bovine serum (FBS) (**Supplemental Table 1B**).

In summary, our experiments demonstrate that GFP-TurboID-ER biotinylates proteins in the secretory pathway of cultured fly cells, which can be recovered from the extracellular space by streptavidin-beads and identified by LC-MS/MS, supporting its utility for *in vivo* tissue-specific secretome profiling.

### Generating and characterizing a collection of transgenic fly lines for in vivo secretome labeling in 10 major tissue types

Following our encouraging results in cell culture, we set out to express TurboID-ER *in vivo* in a tissue-specific manner. To accomplish this, we generated transgenic flies that express GFP-TurboID-ER under the control of the *Gal4*/*UAS* binary expression system^27^ (**Figure 2A**). We constructed 10 fly strains that each carry two transgenes, a *UAS-GFP-TurboID-ER* transgene and a tissue-specific *Gal4* transgene (**Figure 2A**), which results in constitutive tissue-specific expression of TurboID-ER (**Figure 2B**). This approach facilitates raising large numbers of developmentally synchronized flies with the desired genotype. *Gal4* lines were selected to represent most major non-overlapping tissues/organs (e.g. muscle, fat, neurons) and, when possible, express in all cell subtypes (e.g. pan-glial *repo-gal4*). We confirmed tissue-specific expression in each recombinant line by visualizing GFP fluorescence in 3^rd^ instar larvae (**Figure 2C**). Importantly, these 10 lines were viable, fertile, and had no obvious differences in morphology, suggesting constitutive expression of TurboID-ER is non-toxic, like that observed for cytoplasmic TurboID^6^.

**Figure 2:**
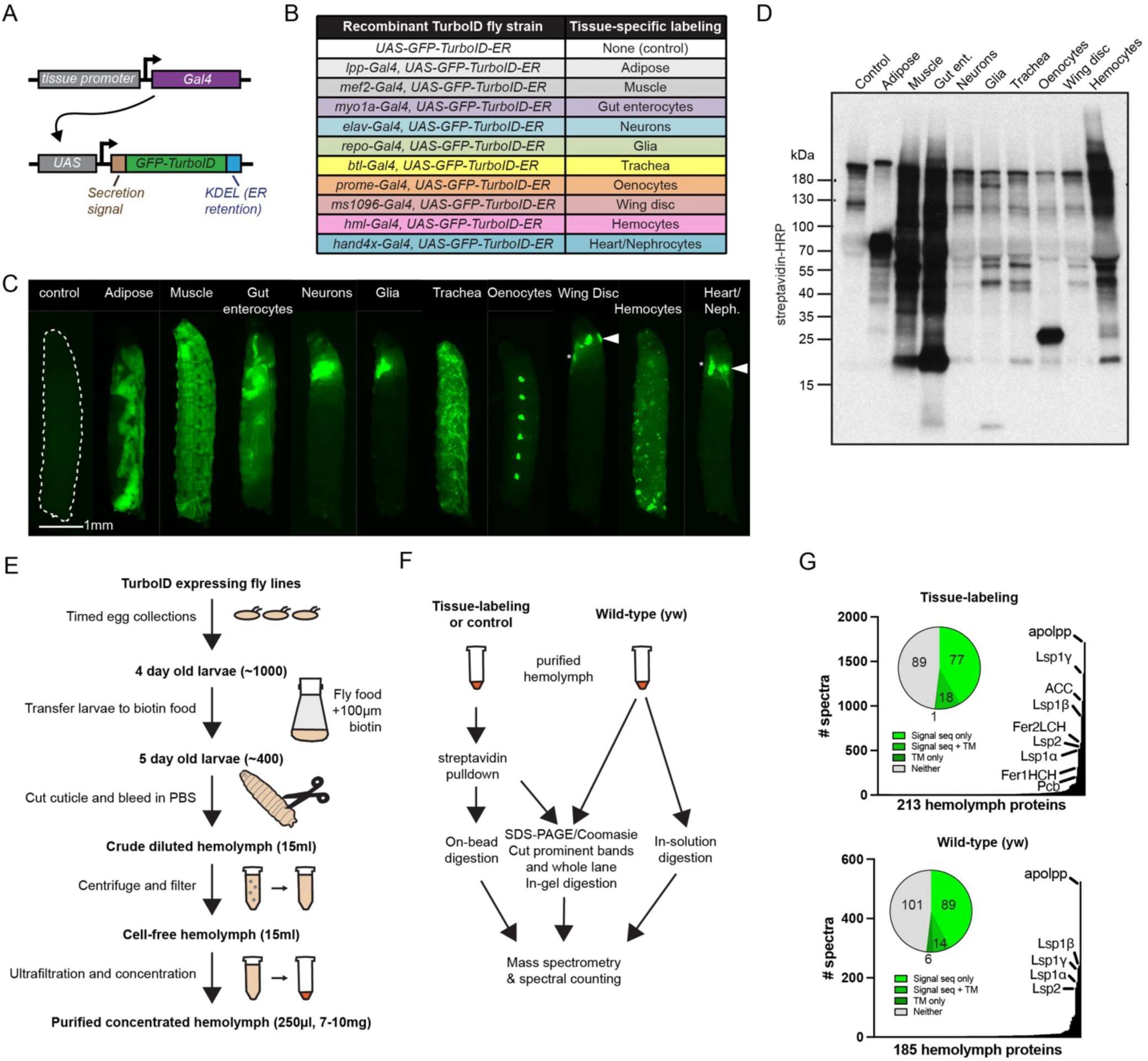
Labeling secreted hemolymph proteins in *Drosophila* 3^rd^ instar larvae using tissue-specific ER-localized TurboID. (A) Schematic of two transgene system to express GFP-TurboID-ER under a tissue-specific promoter (B) Table of recombinant fly strains expressing GFP-TurboID-ER in major tissue types (C) Stereo fluorescence microscopy 3^rd^ instar larvae. Recombinant fly strain expression visualized with *UAS-6x-GFP* crossed to recombinant lines. (D) Western blot of biotinylated proteins in purified hemolymph from 20 3^rd^ instar larvae expressing GFP-TurboID-ER in specific tissues. All lanes loaded 3.6µg protein, except the adipose labeling sample (72ng). (E) Schematic of large-scale hemolymph isolation and purification from 3^rd^ instar larvae. (F) Schematic of mass spectrometry experiments involving purified hemolymph or streptavidin pulldowns. (G) Analysis of proteins identified following mass spectrometry, ranked by # of peptide spectra, and predicted as secreted.

Next, we detected labeled proteins from isolated hemolymph using SDS-PAGE and streptavidin-HRP western blotting. We focused on larval rather than adult hemolymph because it is easier to collect in large quantities and is therefore more amenable to future large-scale experiments (see next results section). We collected hemolymph from 20 wandering 3rd instar larvae per genotype, after raising them on biotin-supplemented food for 24 hours, which we previously showed maximizes biotinylation^6^. As a negative control, we used hemolymph collected from *UAS-GFP-TurboID-ER* larvae (i.e. no Gal4). We observed numerous biotinylated proteins in hemolymph from tissue-specific GFP-TurboID-ER strains (**Figure 2D, Supplemental Figure 2A**). Importantly, the banding pattern and intensity of biotinylated proteins differed among the tissue-specific genotypes. For example, biotinylated proteins appear as a smear of different molecular weights or as discrete protein bands, such as a prominent ∼25 kDa band in hemolymph from oenocyte-labeling (**Figure 2D**). Hemolymph from adipose-labeling gave by far the strongest signal compared to the other samples, agreeing with previous studies that the fat body is a major source of secreted hemolymph proteins^20^. To compensate for this signal disparity, we loaded 1/50th the amount of hemolymph from adipose-labeling in western blots (**Figure 2D, Supplemental Figure 2B**). These results demonstrate that our *in vivo* TurboID-ER labeling approach enables detection of secreted proteins in larval hemolymph and suggests tissue-specific differences in the composition of the secretome.

### A *Drosophila* larval hemolymph collection protocol that is compatible with large-scale streptavidin pulldowns

Our standard hemolymph isolation protocol, which involves bleeding 20 larvae, typically yields 15µg of protein. However, large-scale streptavidin pulldowns for MS require at least 3mg of protein input to ensure sufficient material is recovered after enrichment^28^. In addition, we eventually aimed to perform quantitative MS analysis using biological triplicates for 10 tissue-specific labeling samples. Given our large number of samples and the substantial amount of hemolymph required, it became clear that our standard protocol was not scalable for this purpose. Therefore, we developed a new protocol to isolate large amounts of purified hemolymph from 3^rd^ instar larvae (**Figure 2E**). First, we synchronized development using timed egg collections from each recombinant fly strain expressing *GFP-TurboID-ER*. After 4 days, we transferred larvae to biotin-supplemented food. After 24hrs, we collected ∼400 3^rd^ instar larvae and bled them in PBS. The crude diluted hemolymph was then filtered and centrifuged to remove larvae, tissue debris, and cellular fragments. Finally, we passed filtered cell-free hemolymph through a 3 kDa centrifuge filter to concentrate the sample and remove free biotin, yielding ∼7-10 mg of purified hemolymph (**Figure 2E**).

Next, we performed pilot streptavidin pulldowns and label-free LC-MS/MS using hemolymph isolated with our large-scale procedure. Our goal was to evaluate protein identification using our combined genetic and biochemical strategy, specifically to assess signal quality (i.e. detection of known or predicted secreted proteins), extent of background protein contamination, and the degree of tissue specificity. We focused on tissues that gave strong signal by western blot, including muscle, adipose, gut, glia, and oenocytes, and only used single biological replicates at this stage. We purified hemolymph from larvae with tissue-specific labeling, enriched biotinylated proteins on streptavidin beads, and processed samples for LC-MS/MS by either in-gel (whole lane and prominent Coomassie-stained bands) or on-bead digestion (**Figure 2F, Supplemental Figure 2C**). To help interpret our results, we also collected hemolymph from wild-type larvae and processed it using in-gel digestion of prominent Coomassie-stained bands and in-solution digestion (**Figure 2F, Supplemental Figure 2D**).

In total, we identified 213 proteins from tissue-labeling samples and 185 proteins from wild-type hemolymph samples, with 53% and 52% being predicted secreted, respectively (**Figure 2G, Supplemental Table 2**). The most abundant proteins identified in unlabeled hemolymph are the apolipoprotein Apolpp and the larval serum proteins Lsp1β, Lsp1γ, Lsp1α, Lsp2 (**Figure 2G, Supplemental Table 2, Supplemental Figure 2D**), which is consistent with previous studies^20,29,30^. We readily identified these same abundant proteins in tissue-labeling samples, including other known hemolymph proteins such as ferritins, prophenoloxidases, and extracellular matrix proteins (**Figure 2G, Supplemental Table 2, Supplemental Figure 2C**)^20,31–33^.

Many proteins from tissue-labeling samples exhibited tissue-specific enrichment. For example, Fer1HCH and Fer2LCH, which together form the major ferritin complex and are known to be secreted from the gut^34^, were the top proteins identified from a prominent ∼25 kDa gel band that was specific to gut enterocyte labeling (**Supplemental Figure 2C, Supplemental Table 2**). In addition, Lectin-22c, an uncharacterized C-type lectin protein, was the top protein identified from a prominent ∼35 kDa gel band that was specific to oenocyte labeling (**Supplemental Figure 2C, Supplemental Table 2**). We defined a set of 48 tissue-enriched proteins by comparing the spectral counts of proteins from whole gel lanes (**Supplemental Figure 2E, Supplemental Table 2**). In many cases, identified proteins originated from the expected tissue, such as vkg from fat body^31,35^ and Obp44a from glia^36^ (**Supplemental Figure 2E, Supplemental Table 2**). We also identified known hemolymph proteins from unexpected tissues, such as the adipose proteins apolpp, Lsp1β, and Gel^35,37^ from muscle-labeling samples, and the adipose proteins cv-d, Idgf4, and Tsf1^38–40^ from glia-labeled samples (**Supplemental Figure 2E, Supplemental Table 2**). Excitingly, we identified uncharacterized tissue-specific secreted proteins, such as CG6867 from muscle, CG5080 from glia, Lectin-22c from oenocytes, and CG2233 from adipose (**Supplemental Table 2**). Finally, we identified proteins in tissue-labeling samples that were missing from wild-type hemolymph samples (e.g. lectin-22c, cv-d, bnb, and CG5080) (**Supplemental Table 2**), suggesting that tissue-labeling and pulldown helps enrich low abundance hemolymph proteins.

Our data also highlight unwanted background proteins binding to the streptavidin beads. Two top identified proteins are ACC and Pcb, which are naturally biotinylated proteins^41,42^, and were identified in nearly all streptavidin-enriched samples, but not from wild-type hemolymph (**Supplemental Table 2**). Therefore, ACC and Pcb may be present in circulation, or released from tissue during larval bleeding, and thus bind to streptavidin beads. We also found evidence of non-specific binding of non-biotinylated proteins to the streptavidin beads. For example, the top 5 most abundant proteins in wild-type hemolymph, Apolpp, Lsp1β, Lsp1γ, Lsp1α, and Lsp2 (**Figure 2G**) were identified after streptavidin pulldown from control hemolymph (**Supplemental Table 2**). This phenomenon of non-biotinylated proteins binding to streptavidin beads, despite extensive and harsh bead washes, has been observed by other studies^43,44^.

In summary, our tissue-specific TurboID-ER fly strains effectively label and enrich secreted biotinylated proteins in the hemolymph. These pilot results validate the logic of our approach and provide a strong rationale to scale up for quantitative MS analysis with biological replicates across all 10 tissue types. However, to maximize the impact of these quantitative MS experiments, we first addressed the issue of background proteins binding non-specifically to streptavidin beads.

### Reduced non-specific binding to streptavidin beads with more stringent washing

Non-specific binding of non-biotinylated abundant hemolymph proteins to the beads creates two problems for tissue secretome analysis. 1) It reduces the chances of identifying tissue-secreted biotinylated proteins, particularly those that are low abundance and 2) it prevents a proper analysis of the tissue of origin of abundant hemolymph proteins.

Streptavidin pulldown of wild-type hemolymph revealed a prominent band between 70-100 kDa on SDS-PAGE gels (**Supplemental Figure 2C**), and LC-MS/MS of the entire gel lane identified all four Larval Serum Proteins (LSPs) (**Supplemental Table 2**). Indeed, LC-MS/MS of a similar molecular weight band from wild-type hemolymph (**Supplemental Figure 2D**) revealed that it was predominantly composed of all four Larval Serum Proteins (**Supplemental Table 2**). Therefore, using our current pulldown protocol, which is based on previous studies^6^, we hypothesize that LSPs and other abundant proteins in hemolymph non-specifically bind to streptavidin beads.

To test whether more stringent bead washing could reduce this non-specific binding, we applied a series of harsher wash conditions. 2M Urea is traditionally used as a harsh washing step because it denatures proteins, so we tested increasing Urea concentrations. In addition, others have used SDS in streptavidin bead washing steps^43,45,46^, therefore we tested Urea washes with and without SDS (**Supplemental Figure 3A**). To monitor non-specific binding to the beads, we detected LSPs as a 70-100 kDa protein band on silver-stained gels and by western blotting using anti-LSP-1γ (**Supplemental Figure 3A**), which is cross-reactive with all three LSP-1 proteins^47^. Our results show that higher concentrations of Urea, as well as adding SDS, reduced non-specific binding of LSPs below the level of detection (**Supplemental Figure 3A**).

Next, we tested if more stringent washing had a negative impact on the binding of biotinylated hemolymph proteins to the beads. As described previously (**Figure 2F**), we collected hemolymph from larvae with adipose labeling, performed streptavidin pulldowns, eluted biotinylated proteins, and detected biotinylated proteins on western blots with streptavidin-HRP. Using a panel of Urea and SDS bead washing conditions, there were no obvious differences in the intensity or banding pattern of biotinylated proteins (**Supplemental Figure 3B**). Furthermore, the prominent 70-100 kDa band corresponding to LSPs appears similar in intensity for all bead washing conditions. Therefore, we conclude that harsher bead washing reduces non-specific binding but does not reduce retention of biotinylated proteins on the beads. Thus, we adopted 4M Urea + 2% SDS in future bead washing steps.

### A quantitative proteomic secretome map of 10 major larval tissue types

Building on our pilot LC-MS/MS results in cultured cells and larvae, we created a comprehensive tissue secretome resource from *Drosophila* larval blood, using quantitative proteomics to profile proteins secreted from 10 tissues. Biotin-labeled proteins were enriched from the blood using our improved bead-washing conditions in triplicate, released from streptavidin beads by on-bead digestion, and analyzed by liquid chromatography-tandem mass spectrometry (LC-MS/MS) employing isobaric chemical labeling using TMT reagents for precise relative quantification (**Figure 3A**). A total of 2543 proteins were quantified across the ten experimental groups and fold changes were calculated against enrichments from negative control pulldown samples (**Supplemental Table 3A, Supplemental Figure 4A**). The experimentally enriched proteins showed overrepresentation of proteins containing signal peptide sequences (**Figure 3B**), and principal component analysis showed clear separation of experimental groups from each other and from the control sample (**Supplemental Figure 4B**). To confidently identify candidate secreted proteins, we established fold-change thresholds for each tissue by minimizing nominal false discovery rates using a predefined list of positive-control secreted proteins and negative-control intracellular protein lists (**Supplemental Table 3A**, **Supplemental Figure 4C**, **Methods**). Using these strict thresholds, we report 535 proteins as candidate secreted factors from at least one of these ten tissues (**Supplemental Table 3B**).

**Figure 3:**
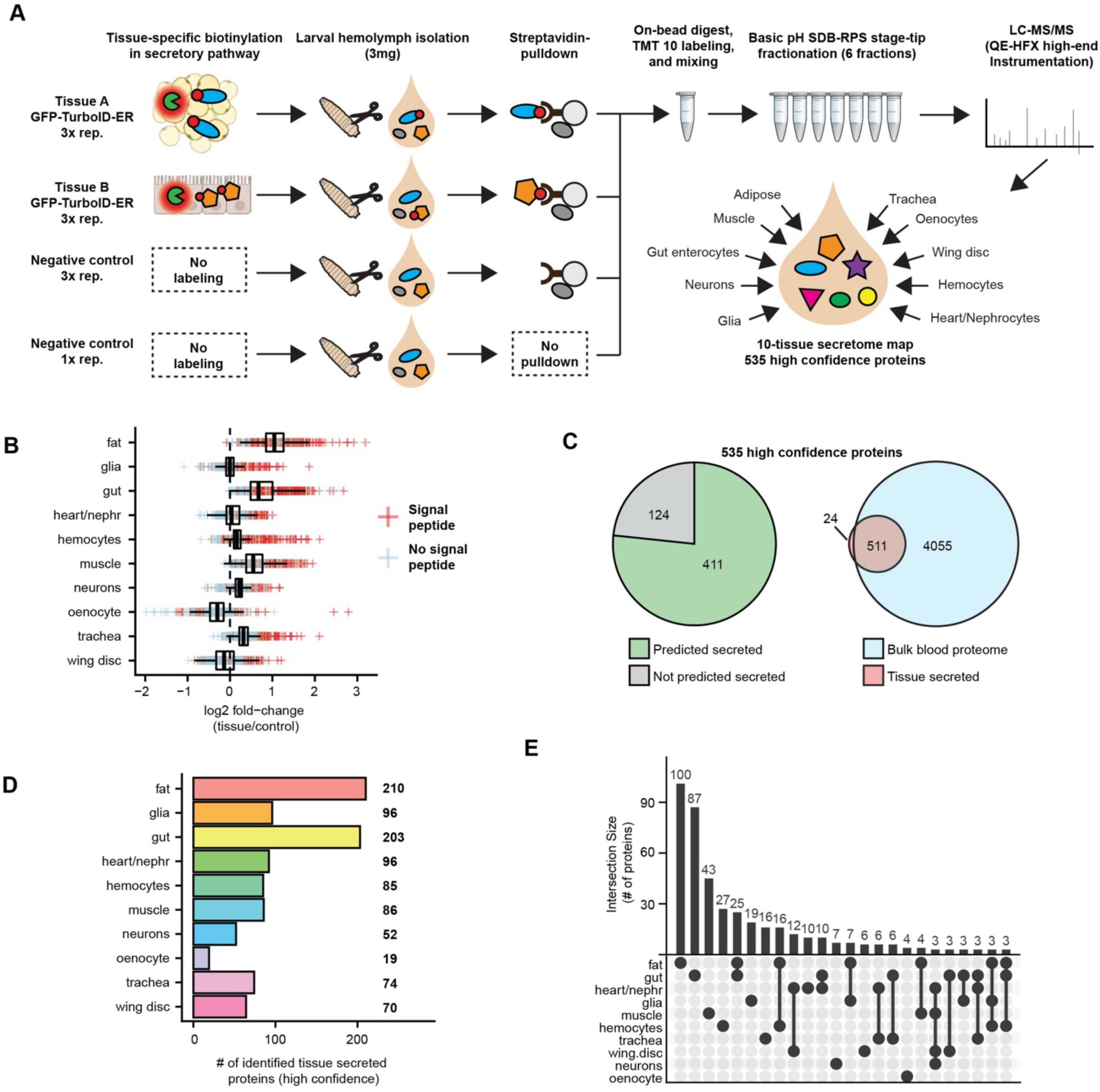
A secretome map of 10 major larval tissues using quantitative proteomics. **(A)** Workflow for proteomic analysis of tissue secreted proteins. **(B)** Boxplots showing log2 fold-change of TMT intensities for each of the 10 tissues relative to the appropriate control. Individual protein values are shown in the plot with color indicating the content of a signal peptide. Enrichment of signal peptide-containing proteins is observed at higher fold-changes for all tissues. Signal peptide annotation by Uniprot. **(C)** Annotation of the 540 proteins secreted by tissues with high confidence. Left pie chart shows number of proteins annotated as possible secreted (Manual literature search, Uniprot database, Flybase database, TMHMM 2.0, SignalP, or DeepLoc) or with no identified secretion annotation. Right venn diagram shows the overlap with proteins identified by proteomic analysis of whole blood from larvae. **(D)** Bar graph showing the number of proteins identified as secreted from each tissue with a high-confidence criteria. **(E)** Upset plot showing the number of proteins identified as secreted by a unique tissue or by multiple tissues. Intersects with less than 3 proteins are not shown. See Supplemental Table 3B.

First, we examined what proportion of the high-confidence 535 proteins are known or predicted to be secreted into hemolymph. Using a combination of protein databases and domain prediction algorithms, we determined that 411 (77%) have the potential to be secreted or shed (**Figure 3C**, **Supplemental Table 3B**). In addition, 267/535 (50%) of proteins are annotated as in hemolymph (Flybase, Uniprot) or present in a previous 3^rd^ instar larval hemolymph dataset^20^ (**Supplemental Table 3B)**. Finally, we generated a label-free 3^rd^ instar hemolymph proteome of 4566 proteins using extensive fractionation for deep-scale analysis (**Supplemental Table 3C**, **Methods**), in which 511 (96%) of our high-confidence proteins were present (**Figure 3C**). In total, 513 (96%) of our high-confidence proteins tissue-secreted proteins are present in at least one of these hemolymph datasets (**Supplemental Table 3B**). This label-free data also highlights the advantages of TurboID-ER labeling, namely tissue-of-origin information, enrichment relative to abundant proteins, and limiting to classically secreted proteins. For example, three top oenocyte enriched proteins (CG7763, Lectin-22C, Lectin-galC1), which are all predicted secreted C-type Lectin proteins, are only ranked #300, #183, and #479, respectively, in label-free data. Furthermore, many cytoplasmic proteins were identified in the hemolymph proteome but not tissue-labeling experiments, such as the muscle-expressed cytoplasmic proteins Mhc, Msp300, and bt (**Supplemental Table 3B-C**).

Among the list of 535 high-confidence secreted proteins are the same major hemolymph proteins seen in our non-quantitative approach (**Supplemental Table 2**), including apolipoproteins, larval serum proteins, ferritins, prophenoloxidases, and extracellular matrix proteins (**Supplemental Table 3B**). We identified additional known hemolymph proteins, such as Myo, Jheh1/2/3, Ance, Idgf2/3, and SDR (**Supplemental Table 3B**)^39,48–51^. Interestingly, many of our high-confidence secreted proteins have not been characterized, including 93 uncharacterized “CG” proteins (**Supplemental Table 3B**).

We also identified peptides corresponding to GFP-TurboID from all 10 tissue-labeling samples (**Supplemental Table 3D)**. This likely corresponds to auto-biotinylated GFP-TurboID that can bind streptavidin beads. Furthermore, GFP-TurboID enrichment was above strict thresholds we used to identify the set of 535 tissue-secreted proteins for 8 out of the 10 tissues. Therefore, we explored the possibility that secreted GFP-TurboID leads to false positive enrichment of hemolymph proteins. Theoretically, if this occurs, then the abundance of proteins in whole hemolymph samples would correlate to those from the streptavidin-enriched samples. Each TMT plex experiment contained raw blood as an internal control in addition to the enrichments from tissue-labeling samples (**Figure 3A, Methods**), and we observed a low correlation between raw blood and enriched samples (**Supplemental Figure 5**). We further collected an independent label-free proteome dataset from 3^rd^ instar hemolymph samples (See Blood Proteome section in Methods). As above, little or no correlation was observed between the abundance of proteins in whole hemolymph samples (label-free dataset) and relative enrichment of proteins from the streptavidin-enriched samples (TMT dataset) (**Supplemental Figure 6** and **Supplemental Table 3E**). Therefore, these data suggest that secreted GFP-TurboID is not labeling blood proteins, or the effect is negligible.

Next, we examined the tissue of origin of the 535 tissue-secreted proteins. The largest number of candidate proteins were observed from the fat and gut with 210 and 203 proteins, respectively, while oenocytes and neurons had the lowest number of candidate proteins with 19 and 52 proteins, respectively (**Figure 3D, Supplemental Table 3B**). Unexpectedly, 319 (59.6%) high-confidence secreted proteins originated from an individual tissue (**Figure 3E, Supplemental Figure 4D, Supplemental Table 3B**). Fat body shared the largest overlap of proteins across other tissues, with 110 identified from fat and at least one other tissue (**Supplemental Table 3B**). Interestingly, by comparing the tissue-enrichment of proteins to their abundance in bulk blood, we find that most proteins with greater tissue-enrichment had middle to low abundances in bulk blood (**Supplemental Figure 6**). The exception was fat body, which is known to secrete extremely abundant proteins such as Apolipoprotein and Larval serum protein. This suggests that tissue-specific ER-labeling and streptavidin enrichment can increase the sensitivity of blood protein identification in addition to determining a tissue of origin. In support of this, we identified 13 predicted secreted proteins using tissue secretome labeling but not from bulk hemolymph (**Figure 3C, Supplemental Table 3B-C**).

To identify possible tissue-specific biological functions in the list of 535 proteins, we performed pathway enrichment analysis (**Supplemental Figure 7A-B, Supplemental Table 3F**). As expected, all the tissues were enriched with proteins containing an “extracellular region” annotation, and most tissues were enriched with “extracellular matrix” annotation. Other pathways shared across many tissues included “defense response” for fat, glia, and hemocytes, and “endopeptidase inhibitor activity” for fat, glia, gut, heart/nephrocytes, and trachea. Some pathways showed unique enrichment to individual tissues, such as imaginal disc growth factor terms for glia and blood pressure related terms for the fat body.

Next, we examined conservation of our list of fly 535 proteins with human. 365 (68%) proteins have at least one human homolog (DIOPT score 3 or higher) (**Supplemental Table 3B**). Furthermore, 232 (43%) and 108 (20%) are homologs of predicted human secreted or plasma proteins, respectively^2^ (**Supplemental Table 3B**). For example, CG6867, CG31999, and CG2493 are homologs of human proteins OLFM4, Fibulin-1, PRCP, respectively, which have also been identified in plasma^52–54^. Furthermore, gut enterocyte-labeled CG9672 is a homolog of human Protein C, which is secreted from intestinal enterocytes and is defective in the inherited blood clotting disorder Protein C Deficiency^55^.

Unexpectedly, 64 of our high-confidence secreted proteins are predicted to have a transmembrane domain (**Supplemental Table 3B**). This includes Sog, which is known to be shed as an ectodomain^56^, as well as Cals, Nep4, and Dg, which have human homologs (Calsyntenin 1/2/3, Neprilysin, Dystroglycan, respectively) that are shed as ectodomains^57^. Indeed, 100% of 184 unique peptides corresponding to 12 single-pass transmembrane proteins map only to the extracellular domain of the protein (**Supplemental Table 3G**), suggesting that at least some of the transmembrane proteins in our list may be released into hemolymph as ectodomains.

### A snRNA-seq atlas of *Drosophila* third instar larvae for mapping tissue-specific secreted proteins

To provide confidence in our dataset, we compared our list of tissue-secreted proteins to publicly available cell-type transcriptomic data of the encoding genes. First, we examined *Drosophila* transcriptomic data, including single cell RNA-seq (scRNA-seq) of the adult fly^58^, larval wing disc^59,60^, larval blood cells^61^, fly kidney^62^, and RNA-seq of dissected larval tissues^63^. These comparisons suggest that 529 (99%) of the genes encoding tissue secreted proteins are expressed in the expected tissue type (**Supplemental Table 4A**). While these data are encouraging, the majority of the transcriptomic datasets used for comparisons are not ideal, due to non-matching developmental timing (adult vs larvae).

To expand on these comparisons, we generated our own single nuclear RNA seq (snRNA-seq) atlas of wandering 3^rd^ instar larvae to match the same developmental timepoint used for the tissue secretome screen (**Figure 4A, Supplemental Table 5A**). Through filtering steps, such correcting for ambient RNA and removing doublets, we identified 28 cell-type clusters at resolution 1.0 (**Figure 4B, Supplemental Table 5B**). Furthermore, we re-clustered eight poorly separated “mixed” clusters into 12 clusters at resolution 0.5 (**Figure 4C, Supplemental Table 5C**). Using a panel of cell-type specific marker genes from the literature and the Fly Cell Atlas as reference^64^, we annotated clusters into broad cell/tissue-types, including those used in our tissue-secretome screen (e.g. muscle, fat body) (**Figure 4B-C**, **Supplemental Table 5B-D, Supplemental Figure 8A**). For example, hemocyte marker genes are enriched in cluster 7, and gut enterocyte marker genes are enriched in cluster 3 (**Supplemental Figure 8B-C**). In addition, using the glial marker gene *repo*, we identified 3,476 putative glial nuclei and computed a list of marker genes (**Supplemental Table 5E**). Similar efforts to identify heart, nephrocyte, and oenocyte nuclei failed, perhaps because these cell-types are less abundant. An interactive version of this 3^rd^ instar larval snRNA-seq map can be found at flyrnai.org.

**Figure 4:**
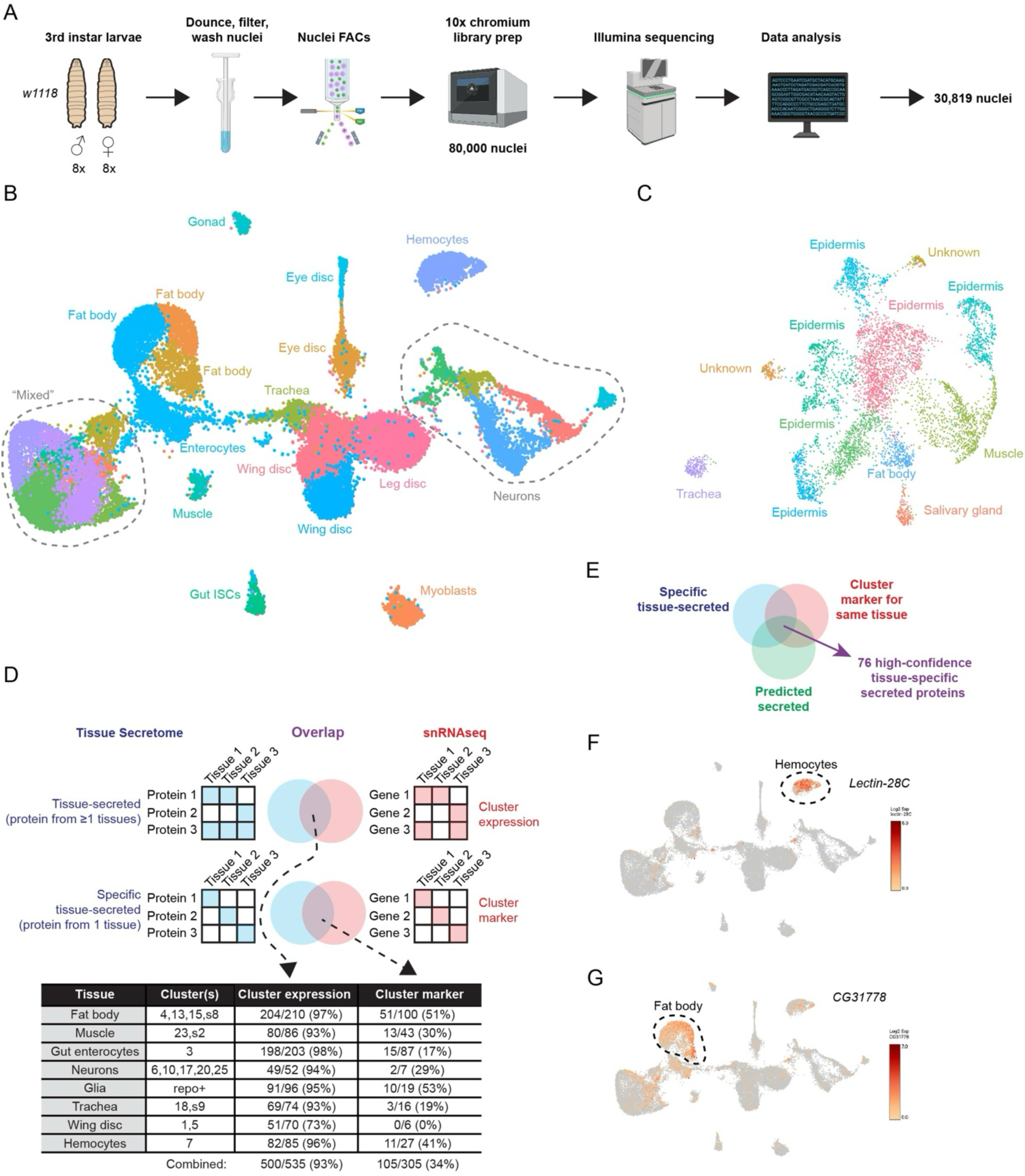
A 3^rd^ instar larval single nuclear RNA-seq map. **(A)** Workflow schematic of larval nuclear isolation, droplet sequencing, and analysis. **(B)** A single UMAP of the 3^rd^ instar larvae containing 28 annotated cell-type clusters, including 8 “mixed” clusters that were re-clustered in **C**. Resolution is 1.0. **(C)** A single re-clustered UMAP of the “mixed” cluster in **B**, annotated as 10 cell-types and 2 unknown cell-types. Resolution is 0.5. **(D)** Strategy to compare tissue secretome proteins with their encoding gene snRNA-seq expression (top). Table with overlap comparisons (bottom). **(E)** Venn diagram showing approach to identify high-confidence tissue-specific secreted proteins. **(F-G)** Depiction of gene expression (heatmap) in UMAP in **B**. Heatmap scale (right) and location of highest cluster gene expression (dotted circle). Genes depicted are **(F)** *Lectin-28c* and **(G)** *CG31778*.

Next, we examined the overlap between our tissue-of-origin proteomic data and tissue-expression of the encoding genes. First, we counted how many genes encoding our 535 tissue-secreted proteins were expressed in the corresponding snRNA-seq cluster(s) (**Figure 4D, Supplemental Table 3B, Supplemental Table 5F-H**). For example, of the 86 muscle-secreted proteins we identified, 80 (93%) of the encoding genes were expressed in the muscle clusters. Second, to measure tissue-specificity, we counted how many genes encoding tissue-specific secreted proteins are marker genes for the same cluster (**Figure 4D, Supplemental Table 3B, Supplemental Table 5B-C, E**). For example, of the 43 muscle-specific-secreted proteins identified by LC-MS/MS, 13 (30%) of the encoding genes were muscle cluster marker genes (**Figure 4D, Supplemental Table 3B, Supplemental Table 5B-C, F-G**). In total, these comparisons for 8 tissue types show that 93% (500 out of 535) of tissue-secreted proteins are expressed in at least one corresponding tissue cluster(s) and 34% (105 out of 305) of genes encoding tissue-specific-secreted proteins are marker genes for the expected cluster(s) (**Figure 4D**).

Finally, we defined a list of 76 high-confidence tissue-specific secreted proteins as 1) tissue-specific proteins from our proteomic screen, 2) encoded by marker genes for the same tissue cluster(s), and 3) predicted to be secreted or transmembrane (**Figure 4E, Supplemental Table 5K, Supplemental Figure 9**). In this list, we found known tissue-specific hemolymph proteins, such as Myo from muscle, NimB1 from hemocytes, and ADGF-D from fat body (**Supplemental Table 5K**). Interestingly, this list contains mostly uncharacterized proteins, such as Lectin-28C and CG31778, which are enriched in hemocytes and fat body, respectively (**Figure 4F-G, Supplemental Figure 9**).

### A collection of knock-in flies to characterize tissue-specific secreted proteins *in vivo*

Next, we used *in vivo* knock-in fly strains to test if proteins from our proteomic screen are expressed in the expected tissue or are present in circulation (**Figure 5A**). We chose to target 22 proteins based on their tissue specific expression, lack of characterization, presence in the blood, and prediction as secreted/transmembrane (**Supplemental Table 6**). Included in this list are seven C-type lectin proteins (e.g. lectin-22C), six proteases/protease-inhibitors (e.g. CG31821/SCPEP1), two proteins with no predicted functional domains or characterization (CG3777, CG9917), and a homolog of vertebrate olfactomedin proteins (CG6867).

**Figure 5:**
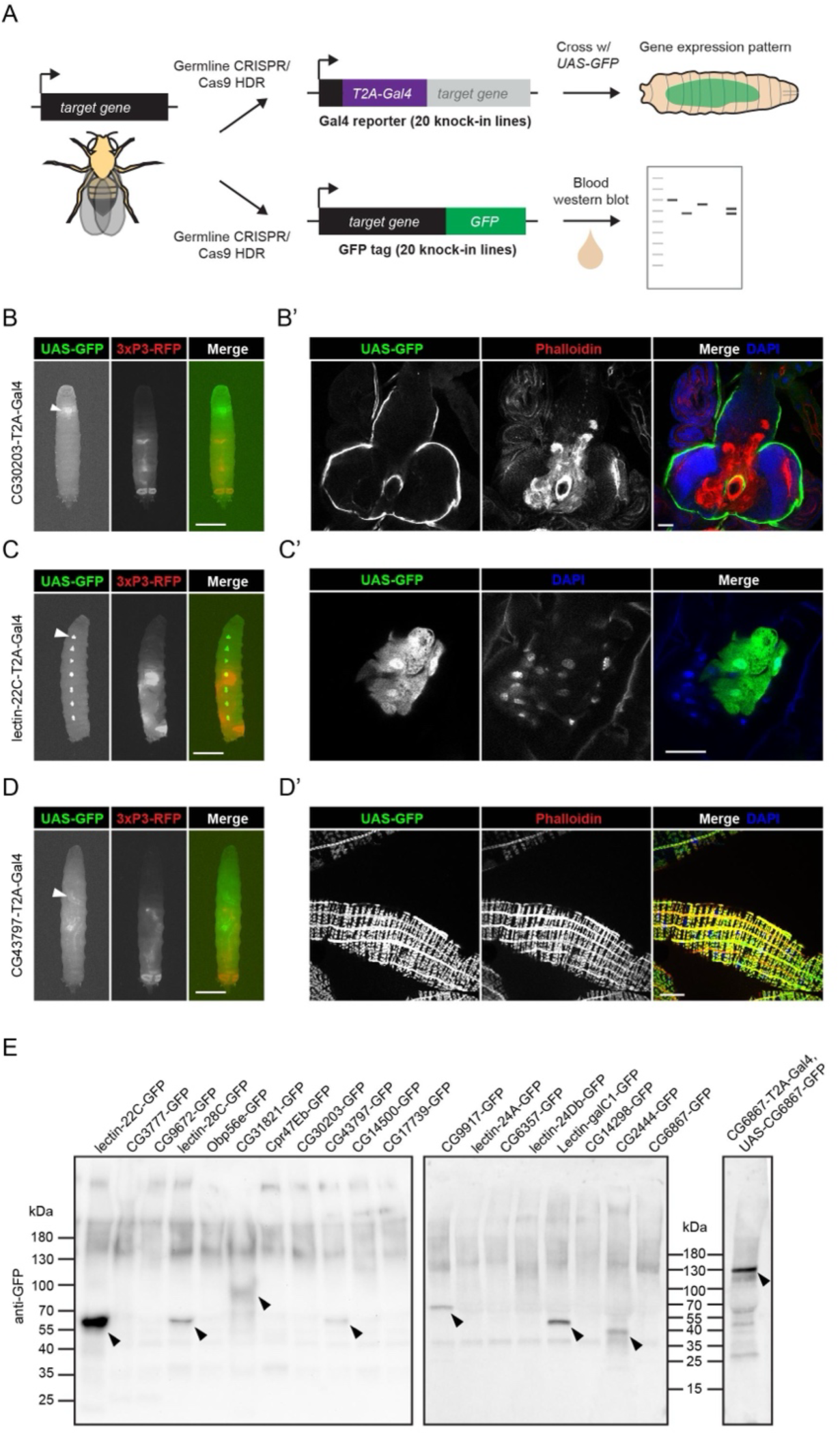
In vivo characterization of uncharacterized hemolymph proteins using knock-in fly lines. (A) Schematic of generating fly knock-ins and experimental use to assay gene expression and protein localization. **(B-D)** Widefield fluorescence microscopy images of whole 3^rd^ instar larvae expressing *UAS-GFP* (green) under the control of the *gene-T2A-Gal4* transgene. Also shown is *3xP3-RFP* (red) marker fluorescence. Arrowheads indicate localized GFP fluorescence in the **(B)** brain, **(C)** oenocytes, and **(D)** visceral gut muscle. Scale bar is 1mm. **(B’-D’)** Confocal microscopy images of dissected tissues from 3^rd^ instar larvae, showing anti-GFP (green), phalloidin-660 (red), and DAPI (blue) stains. Scale bars are 50µm. **(B-B’)** *CG30203-T2A-Gal4*, **(C-C’)** *lectin-22C-T2A-Gal4*, **(D-D’)** *CG43797-T2A-Gal4*. **B’** and **C’** are confocal slices, **D’** is a projection. **(E)** Western blot of 3^rd^ instar larval hemolymph collected from GFP-tag lines. Negative controls are intracellular GFP-tagged proteins and positive controls are GFP-tagged known hemolymph proteins. Arrowheads indicate bands at the expected molecular weight. (58.8 kDa = Lectin-22c-GFP; 59.2 kDa = Lectin-28c-GFP; 77.4kDa = CG31821-GFP; 56.2 kDa = CG43797-GFP; 63.1 kDa = CG9917-GFP; 50.8 kDa = Lectin-galC1-GFP; 41.4 kDa = CG2444-GFP; 134.3 kDa = CG6867-GFP)

We generated knock-in fly strains using CRISPR/Cas9 and homology directed repair (HDR) in the fly germline as previously described^65^. For gene expression analysis, we inserted a *T2A-Gal4* reporter gene^66^ into 5’ coding sequence (**Figure 5A**). These *T2A-Gal4* reporter lines are also likely null alleles since they are predicted to truncate the wild-type protein product. To detect the corresponding target protein, we inserted *GFP* into 3’ coding sequence to create a C-terminal GFP tag (**Figure 5A**). To facilitate the efficient isolation of these knock-in lines, we developed modified donor plasmids that allow users to select against off-target plasmid integration (**Methods**), which will be described in a separate study.

Gene expression patterns were visualized by crossing *T2A-Gal4* lines to *UAS-GFP* flies and imaging GFP fluorescence in 3^rd^ instar larvae (**Figure 5A**). 19 out of 20 lines exhibited expression in the expected tissue(s) (**Supplemental Table 6, Figure 5B-D, Supplemental Figure 10**). This includes cases where Gal4 expression closely mirrors that seen in our tissue-secretome data, such as *CG30203-T2A-Gal4*, *Lectin-22c-T2A-* Gal4, and *CG43797-T2A-Gal4*, which are specifically expressed in perineural glia, oenocytes, and gut visceral muscle, respectively (**Figure 5B-D, Supplemental Table 6**). One line did not match our TurboID labeling experiments - *Obp56e-T2A-Gal4* is expressed primarily in tracheal spiracles, however Obp56e was enriched from heart/nephrocyte and wing disc (**Supplemental Figure 10**). Since *Obp56e* transcript is not detected in wing discs^60^ (**Supplemental Table 5F**) or larval nephrocytes^67^, we believe that this is a false positive. One possibility is that the wing disc Gal4 driver (*MS1096-Gal4*) expresses in tracheal spiracles.

Next, we determined if GFP-tagged target proteins were circulating in hemolymph. We collected hemolymph from 3^rd^ instar larvae and performed anti-GFP western blotting. Encouragingly, we detected a band at the expected molecular weight for 7 out of 19 GFP-tagged lines (**Figure 5E**). Only 2 out of 5 positive control GFP-tagged proteins were detected using this method (**Supplemental Figure 11**), indicating a high false-negative rate and likely explaining why the remaining 11 GFP-tagged proteins were not detected in hemolymph.

Finally, we asked whether any target proteins localize to tissues different from their site of synthesis, suggesting potential inter-organ functions. For CG2444, gene expression was restricted to the fat body, yet, remarkably, the protein localized to cuticle (**Figure 6A**), an important extracellular matrix on the surface of the animal. Similarly, CG6867 was expressed in somatic muscle, and the protein accumulated in imaginal discs (**Figure 6B**), which develop into adult structures such as the wing. Both CG2444-GFP and CG6867-GFP were detected in hemolymph (**Figure 5E**), supporting their secretion and circulation via the blood. Notably, for CG6867, we only detected the protein on imaginal discs and in the blood using a UAS-CG6867-GFP transgene and not with the endogenous CG6867-GFP knock-in line, suggesting the endogenous signal may be below the detection limit. Together, these results suggest that CG2444 is trafficked from fat-to-cuticle, and CG6867 is trafficked from muscle-to-imaginal disc (**Figure 6C**).

**Figure 6:**
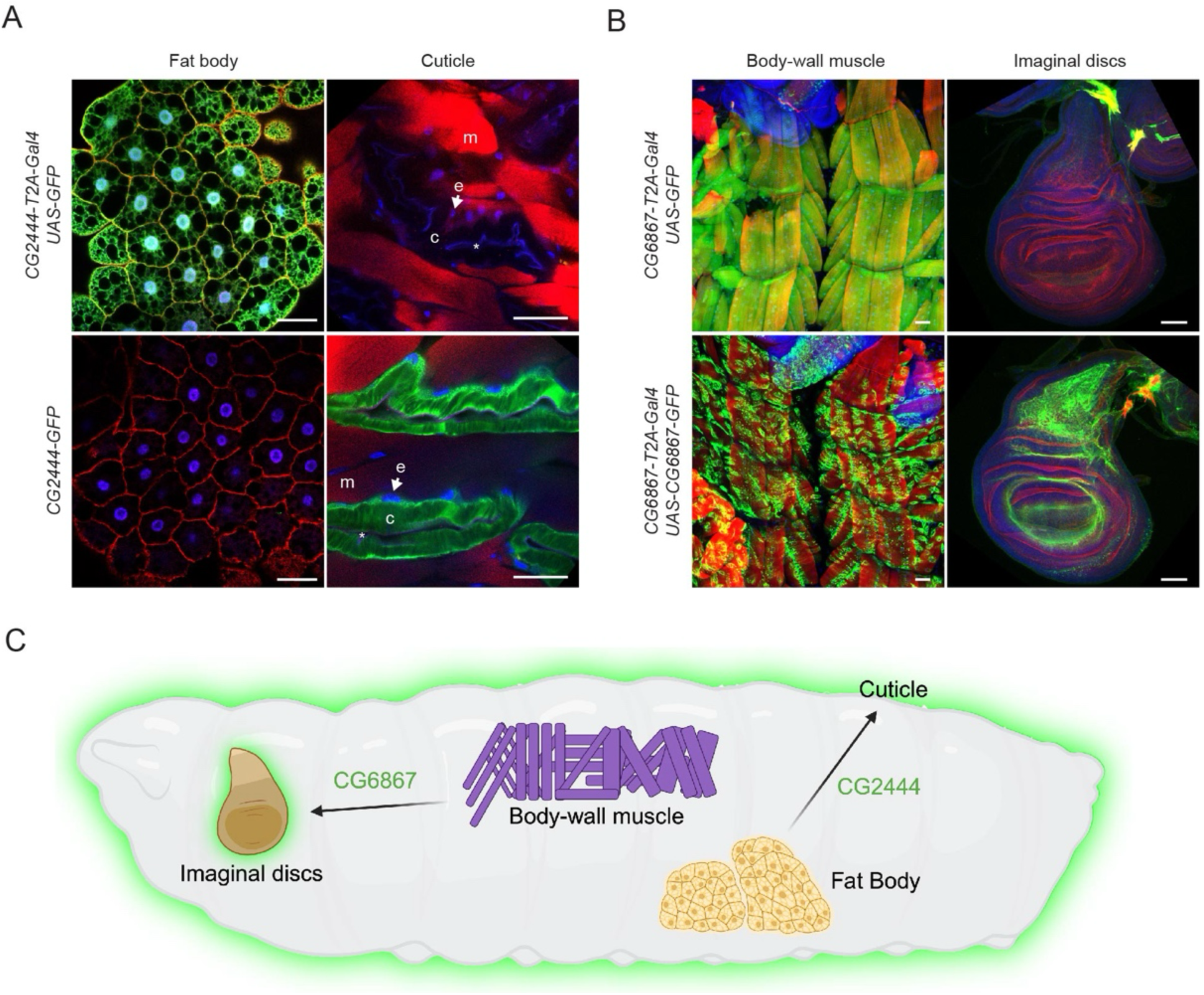
Inter-organ localization of hemolymph proteins. **(A-B)** Confocal microscopy images of dissected tissues from 3^rd^ instar larvae with indicated genotypes, showing anti-GFP (green), phalloidin (red), and DAPI (blue) stains. Images of the same tissue were acquired using identical confocal settings to allow direct comparison of fluorescence intensity. Scale bars are 50µm. **(A)** Images are confocal slices. m = muscle, e = epidermal cell nuclei, c = cuticle, asterix (*) = outside face of cuticle. **(B)** Images are confocal projections. **(C)** Schematic summary of inter-organ proteins synthesized in source tissues and localized to distant recipient tissues.

In summary, our knock-in fly strains validate our tissue-secretome dataset as biologically accurate and containing tissue-specific hemolymph proteins and candidate mediators of inter-organ communication.

## Discussion

This study presents a comprehensive *in vivo* tissue-secretome map in *Drosophila melanogaster* by integrating proximity labeling, mass spectrometry, single-nucleus RNA-seq, and CRISPR knock-in technology. We report new genetic tools, protocols, and datasets that will broadly benefit the scientific community working on secreted proteins and beyond. For example, our tissue-secretome map and *in vivo* characterization enables a simple query of secreted proteins and cell-types of interest, can help assign functions to uncharacterized tissue-specific secreted proteins, and may be a source of human-fly conserved tissue biomarkers.

We report new cell lines and fly strains for labeling secretory proteins in *Drosophila*. In particular, our *UAS-GFP-TurboID-ER* fly lines (insertions on either chromosome II or III) are compatible with thousands of Gal4 lines^68^ to label secreted proteins in a cell-type-specific manner. Furthermore, the generation of *Gal4/UAS-GFP-TurboID-ER* recombinant lines is made simple by screening for GFP fluorescence. We provide a set of 10 premade two-transgene *Gal4/UAS-GFP-TurboID-ER* lines in this study, which express in major tissue types (e.g. muscle, glia, neurons, etc.) and should be useful to the *Drosophila* community to identify secreted proteins or intracellular ER/Golgi-resident proteins.

In addition, we report new protocols for isolating *Drosophila* hemolymph and optimizing streptavidin pulldowns. Previous hemolymph collection methods from larvae or adults were not scalable for large-scale streptavidin pulldowns and often do not remove hemocytes and cell debris^20,69,70^. Our protocol yields >3mg of soluble hemolymph proteins in ∼30 minutes, which is comparable to a human blood plasma sample^71^ and is sufficient for streptavidin pulldown and LC-MS/MS^28^. We used larvae because they yield large amounts of hemolymph^72–74^, but this protocol could in theory be adapted for adult flies. Similarly, while we focused on soluble proteins, this protocol could be applied to circulating exosomes, metabolites, and extracellular RNA. Finally, we empirically determined a wash buffer (2% SDS and 4M Urea) that reduces non-specific binding of abundant LSP proteins to streptavidin beads. While it is unclear whether LSP proteins are representative of other non-specific binders, the inclusion of SDS and high urea concentrations is likely beneficial. Similar conditions have been used by others to reduce background binding to streptavidin beads^43,45,46,75^.

We report new proteomic datasets of secreted proteins in *Drosophila*. Using TurboID-ER labeling, we identified 72 proteins secreted from S2R+ cells into cell culture media. To our knowledge, this is the first secretome for any *Drosophila* cell line, and it highlights uncharacterized proteins such as CG5080. Most importantly, we report an *in vivo* tissue-secretome map of 535 proteins in *Drosophila* hemolymph from 10 major tissue types using TurboID-ER labeling. While other studies used *in vivo* TurboID-ER labeling in mice^8,11–16,18^, our study expands the number of tissues to provide a more comprehensive tissue secretome. While a previous study used BirA*-G3-ER to identify proteins trafficking between tissues in *Drosophila*^18^, our study focused on circulating proteins, offering a more scalable approach using hemolymph-only collection and enabling direct cross-sample comparison. Finally, using label-free MS of larval hemolymph, we identified 4,566 proteins, which is four times more than a previous study^20^. Together, these proteomic resources represent a new comprehensive dataset of *Drosophila* secreted proteins.

We report a snRNA-seq atlas of whole third instar *Drosophila* larvae, which we used to compare our proteomics results. While previous studies profiled dissected larval tissues^59–61^, this is the first snRNA-seq dataset from intact whole larvae, and it is publicly accessible at flyrnai.org. Interestingly, our snRNA-seq dataset revealed at least two clusters of unknown identity. We identified cell-types corresponding to 8/10 tissues used for TurboID labeling and found convincing overlap between the genes expressed as well as tissue-specific expression. We did not identify oenocytes and heart/nephrocytes, perhaps because our dataset was relatively small (∼30k nuclei) and these cell-types are less frequent. In addition, not all proteins identified as tissue-specific by TurboID-ER labeling corresponded to cluster-specific marker genes in the snRNA-seq data, which may reflect post-transcriptional regulation, regulated secretion, or limitations in the sensitivity of either method.

We also report a collection of 40 CRISPR/Cas9 knock-in fly strains, which we used to investigate the quality of our proteomics data. All but one Gal4 line expressed in the expected tissue(s), including remarkably specific patterns in oenocytes, fat body, and hemocytes, as well as sub-cell-types within tissues, such as perineural glia and enterocytes in a sub-region of the midgut. These new Gal4 lines may also be valuable tools for gene-function studies, including expression drivers for circular gut muscles and mouth hook muscles (which to our knowledge are not represented by existing Gal4 lines), as well as loss-of-function alleles. Many of our GFP knock-in lines showed GFP signal in hemolymph or localized to distant tissues, confirming their secretion. Because we generated knock-in fly strains for largely uncharacterized genes, this *in vivo* data opens new areas of investigation into the functions of these proteins. For example, *CG31821-T2A-Gal4* is homozygous lethal, suggesting this gene is essential, and may have functions related to visceral muscle like its human homolog SCPEP1 which is expressed in smooth muscle^76^ and present in plasma^1^. Finally, we showed tissue-specific expression of seven previously uncharacterized C-type lectins (e.g. Lectin-22c) in the gut, muscle, oenocytes, fat body, and hemocytes. Given the established roles of C-type lectins in innate immunity, their distinct expression patterns may indicate coordinated immune functions across multiple tissues.

From our tissue secretome map, we identified two novel proteins that localize to a different tissue than they are expressed, suggesting they have inter-organ functions. CG2444 belongs to the single von Willebrand factor C-domain (SVC proteins) family^77^, which are thought to respond to environmental challenges. Since the fat body is known to be an integrator of metabolic and immune changes^78^, and the cuticle is a barrier to the environment including pathogens^79^, CG2444 may be remote factor that the fat body uses to alter cuticle composition or function. Interestingly, our results suggest a surprising mechanism of CG2444-GFP transcytosis through epidermal cells, like that observed for Serp transcytosis through tracheal cells^80^. CG6867 is the single fly ortholog of the vertebrate olfactomedin family and closely resembles the human transmembrane protein Gliomedin^81^. Since the ectodomain of Gliomedin is known to be shed^82^ and binds to the extracellular matrix (ECM) protein Perlecan^83^, we hypothesize that CG6867 is a muscle-secreted protein that integrates into the ECM of distal tissues such as the imaginal discs. Future studies of these candidate inter-organ factors should investigate their roles in target tissues. Our discovery of these factors demonstrates that our screen, although designed to identify the origins of secreted proteins, can also uncover proteins involved in inter-organ communication

An important concern with TurboID-ER is that the enzyme itself may be secreted and label proteins extracellularly, thus confounding the tissue-specificity imparted by its genetic expression. We did detect GFP-TurboID-KDEL in S2R+ media and in larval hemolymph, likely due to overwhelming the KDEL receptor or cell culture artifacts like cell lysis. This concern might be addressed by reducing expression of TurboID-ER. However, several lines of evidence suggest that extracellular labeling is minimal and does not confound our results. First, ATP levels are low in the extracellular space^84^ and extracellular labeling with TurboID requires added ATP^85^. Second, we see minimal labeling of abundant cow proteins (from fetal bovine serum) in S2R+ cell culture media, which are not visible on stained protein gels but are detectable by MS. Third, we see clear tissue-specific labeling *in vivo* rather than widespread labeling of abundant proteins, and the proteomic profiles from TurboID pulldown and total hemolymph were largely distinct. Finally, this approach of appending a KDEL sequence to TurboID or ER retention has been used by others to label tissue-specific secreted proteins^8,11,13–16,18^. Together, these results indicate that while some TurboID-ER is secreted, it does not significantly impact the specificity of our secretome mapping strategy.

Another concern is that our identification of intracellular proteins in the extracellular space may reflect artifacts caused by TurboID labeling. For example, our list of 535 tissue-secreted proteins contains a small number of ER-resident, cytoplasmic, and transmembrane proteins. TurboID-ER expression could in theory cause biotinylated proteins to mislocalize, such as by protein misfolding, ER-stress, or saturation of KDEL receptors. However, 96% of the proteins identified in our tissue-secretome map were also present in hemolymph from wild-type animals, and our fly strains constitutively expressing TurboID-ER were viable and fertile, arguing against widespread mislocalization or ER-stress toxicity. Furthermore, GFP-tagged knock-in lines confirmed secretion into the hemolymph for many of these proteins. We also speculate that labeled cytoplasmic proteins in the extracellular space may arise from mislocalization of TurboID-ER and subsequent cell leakage, either of which could be tissue-specific. Indeed, we identified seven ribosomal proteins (e.g. RpL8) specifically from muscle. Finally, transmembrane proteins likely represent shed ectodomains rather than intact membrane proteins, as supported by peptide mapping and their lack of exosome annotations. Therefore, we argue that these possible artifacts do not impede us from identifying true secreted proteins.

We note important limitations with our *in vivo* TurboID-labeling strategy to identify secreted proteins. For example, TurboID-ER labels surface-exposed lysine residues on proteins in the secretory pathway, and LC-MS/MS may miss low abundance labeled proteins. Indeed, we did not identify enolase or nplp2 in our tissue-secretome map, despite these being abundant secreted proteins in larval hemolymph^20^ (Supplemental Table 3C) that are each expressed in fat body^86,87^ (Supplemental Table 5B). Similarly, many hormones such as *Drosophila* insulin-like peptides (Dilp1-8) and Unpaireds (Upd1-3) were absent, likely due to their small size and/or low abundance. We did identify the hormone Myoglianin, suggesting the sensitivity threshold lies somewhere between these proteins. Our strategy will likely not identify extracellular proteins secreted via unconventional pathways, such as those bypassing the ER or released in vesicles like exosomes, since TurboID-ER only labels proteins that transit through the ER. Finally, the strength and tissue-specificity of Gal4 drivers can affect whether proteins are identified. Some Gal4 drivers exhibit expression outside the desired domain, such as *repo-Gal4* (glial) and *MS1096-Gal4* (wing disc) that also express in salivary glands. Conversely, some drivers may be too spatially restricted, such as *MS1096-Gal4*, which is limited to the wing disc pouch.

Our datasets provide a foundation for further exploration and open new avenues to generate datasets with even greater resolution and utility. Our study enables queries for specific fly proteins or tissues, and we have already identified at least two candidate inter-organ proteins, highlighting is potential to discover new biology. Additionally, this resource may facilitate the discovery of human biomarkers, as illustrated by our identification of over 100 fly proteins whose human homologs are secreted or detected in plasma. Future studies employing additional Gal4 drivers could provide a more comprehensive tissue secretome by targeting tissues not examined here, such as the ring gland or epidermis. Furthermore, this method, initially established under wild-type conditions in this study, could be applied to disease models to identify tissue-specific induced factors that promote or prevent pathology. Finally, our large-scale hemolymph collection and tissue-specific TurboID-ER–based labeling approaches could be adapted to other insect species, both *in vivo* and in cell culture, broadening their impact across species including disease vectors.

## Acknowledgements

We thank Illia Droujinine and Chiao-Lin Chen for advice on proximity labeling, Jean-Antoine Lepesant for kindly providing anti-LSP-1γ antibodies, Tzu-Chiao and Hongjie Li for advice with snRNA-seq, Raghuvir Viswanatha and Enzo Mameli for assistance with FACs single nuclei sorting, Arpan Ghosh for advice on hemolymph collection, Rich Binari for general lab assistance, and Carlie Widdison, Dhruti Patel, Mia Krause, Sofia Moretti, Jenna Larsen, and Ruben Andres for maintaining fly stocks. We also thank Andy McMahon and his lab for useful discussions. bioRender was used to make a subset of figure panels. JAB was supported by the Damon Runyon Foundation and NIH T32 GM007748/GM/NIGMS. NP was supported by NIH grants R01GM084947, R01GM067761, R24OD019847. PMJB was supported by NIH grant F32HL154711. NP is an investigator of the Howard Hughes Medical Institute.

This article is subject to HHMI’s Open Access to Publications policy. HHMI lab heads have previously granted a nonexclusive CC BY 4.0 license to the public and a sublicensable license to HHMI in their research articles. Pursuant to those licenses, the author-accepted manuscript of this article can be made freely available under a CC BY 4.0 license immediately upon publication.

## Figure Legends

Supplemental Table 1: S2R+ secretome LC-MS/MS results and annotations

Supplemental Table 2: Non-quantitative *in vivo* tissue secretome LC-MS/MS results and annotations

Supplemental Table 3: Quantitative *in vivo* tissue secretome LC-MS/MS results and annotations

Supplemental Table 4: Comparison of tissue secretome to publicly available *Drosophila* and human public RNA-seq datasets

Supplemental Table 5: Comparison of tissue secretome to 3^rd^ instar snRNA-seq Supplemental Table 6: Fly CRISPR knock-in information

Supplemental Table 7: Oligos, dsDNA, and gateway plasmids

Supplemental File 1: Amino acid sequence of GFP-TurboID-ER

## Methods

### Plasmid cloning

Plasmid DNAs were constructed and propagated using standard protocols. Briefly, chemically competent TOP10 E. coli. (Invitrogen, C404010) were transformed with plasmids containing either Ampicillin or Kanamycin resistance genes and were selected on LB-Agar plates with 100 µg/ml Ampicillin or 50 µg/ml Kanamycin. Similarly, gateway destination plasmids containing the *ccdb* toxic gene were propagated in *ccdb*-resistant E. coli (Invitrogen, A10460) by culturing in LB containing 100 µg/ml Ampicillin and 25 µg/ml Chloramphenicol. Oligo and dsDNA sequences are in **Supplementary Table 7A**.

*pMT-EGFP-V5* (Addgene #240240, DGRC #1696) was generated by EcoRI/XbaI (NEB, R3101S, R0145S) restriction enzyme digestion and ligation of PCR-amplified *EGFP* sequence into *pMT/V5-His B* (Invitrogen, V4120-20).

#### Entry vectors

Construction of pEntr vectors (for Gateway cloning) was performed by Gibson assembly (NEB, E2621S) of PCR amplified backbone from *pEntr-dTOPO* (Invitrogen C4040-10) and PCR amplified gene coding sequence (when appropriate, with or without stop codon).

*pEntr_sfGFP-TurboID* (Addgene #240226, DGRC #1682) and *pEntr_BiP-sfGFP-TurboID-KDEL* (Addgene #240223, DGRC #1679) were generated by Gibson assembly of overlapping PCR-amplified fragments containing *sfGFP*, *TurboID*, and *pEntr* backbone sequences. *sfGFP* (superfolder GFP) was amplified from *pCRISPaint-sfGFP-3xP3-RFP* (Addgene #127566)^65^. *TurboID* was amplified from *pUAS-V5-TurboID-NES* (Addgene #116904)^6^. *BiP* and *KDEL* sequences were added by oligos.

*pEntr_HA-spGFP11* (Addgene #240224, DGRC #1680) was generated by Gibson assembly of a *HA-spGFP11* gBlock DNA fragment into *pEntr* backbone sequence.

*pEntr_spatzle-HA* (Addgene #240227, DGRC #1683) and *pEntr_hhN-HA* (Addgene #240225, DGRC #1681) were generated by Gibson assembly of overlapping PCR-amplified fragments containing *spatzle* or *hhN*, and *pEntr_HA-spGFP11* backbone sequences. *spatzle* sequence was amplified from larval cDNA. *hhN* was amplified from *Actin Hh-FL*^88^. hhN is an N-terminal portion of Hh that normally results after protein cleavage. Full length spatzle can be cleaved by Easter^89^, however this is unlikely to occur in S2R+ cells which express low levels of Easter (DGET, https://www.flyrnai.org/tools/dget/web/). In both cases, the C-terminal HA tag is fused to the mature signaling portion of the protein.

*pEntr_CG6867_nostop* (Addgene #240238, DGRC #1694) was generated by Gibson assembly of PCR-amplified coding sequence into *pEntr* backbone sequence. *CG6867* was amplified from cDNA clone FI21454 (DGRC #1661063).

#### Gateway vectors

*pWal10-roe-sfGFP* (Addgene #240239, DGRC #1695) was generated by PCR amplification of *sfGFP* sequence from *pCRISPaint-sfGFP-3xP3-RFP* (Addgene #127566)^65^ and ligation into *pWal10-roe*^90^ that was digested with XbaI (NEB, R0145).

#### Gateway cloning LR reactions

*pMT-sfGFP-TurboID* (Addgene #240231, DGRC #1687)*, pMT-sfGFP-TurboID-ER* (Addgene #240230, DGRC #1686)*, pUAS-sfGFP-TurboID-ER* (Addgene #240232, DGRC #1688)*, pMT-Myc-BirA*G3* (Addgene #240233, DGRC #1689)*, pMT-Myc-BirA*G3-ER* (Addgene #240234, DGRC #1690)*, pUAS-spatzle-HA* (Addgene #240235, DGRC #1691)*, pUAS-hhN-HA* (Addgene #240236, DGRC #1692), and *pUAS-CG6867-sfGFP* (Addgene #240237, DGRC #1693) were constructed by Gateway cloning. Cloning reactions were performed using LR Clonase II Enzyme mix (Invitrogen 11791– 020). Published entry plasmids used were *pEntr_Myc-BirA*-G3* and *pEntr_Myc-BirA*G3-ER* (Addgene # 173814)^18^. Published destination plasmids used were *pWalium10-roe*^90^ and *pMK33-GW*^91^. See **Supplementary Table 7B** for information on assembled gateway plasmids.

#### Scarless GFP and Gal4 HDR donor plasmids with a GMReya(shRNA) marker

*pHD-sfGFP-ScarlessDsRed-GMR-eya(shRNA)* (Addgene #240228, DGRC #1684) and *pHD-T2A-Gal4-3xP3DsRed-GMR-eya(shRNA)* (Addgene #240229, DGRC #1685) were cloned by first mutating the EcoRI site in *pBS-GMR-eya(shRNA)* (Addgene #157991)^92^ using site directed mutagenesis (NEB, E0554S) to produce *pBS-GMR-eya(shRNA)_EcoRImut* (Addgene #240222, DGRC #1678). *sfGFP-ScarlessDsRed* was PCR amplified from *pHD-sfGFP-ScarlessDsRed* (Addgene # 80811)^93,94^, and *T2A-Gal4-ScarlessDsRed* was PCR amplified from Bosch CRISPaint plasmid *pCRISPaint-T2A-GAL4-Hsp70*^95^. Each fragment was cloned into *pBS-GMR-eya(shRNA)_EcoRImut*, which had been digested SacI and HindIII (NEB, R3156S, R3104S), by Gibson assembly.

#### Cloning homology arms into donor plasmids

For T2A-Gal4 knock-ins, 1-2kb genomic homology arms were designed that flank a sgRNA target cut site in 5’ coding sequence. Homology arms were PCR amplified from *nos-Cas9* fly genomic DNA, such that genes on Chr. X or II were amplified from *nos-Cas9attP2*, and genes on Chr. III were amplified from *nos-Cas9attP40*.

For sfGFP knock-ins, 1-2kb genomic homology arms were designed that flank a sgRNA target cut site near the start (for N-terminal fusion) or stop codon (for C-terminal fusion). Homology arms were amplified as for T2A-Gal4 knock-ins. In addition, we introduced 4-5 mutations in the PAM and spacer sequence that introduce synonymous codons in coding sequence. Mutations were encoded on PCR primer 5’ extensions or a synthesized dsDNA fragment that bridged the wild-type homology arm and the insert.

PCR amplified homology arms, and in some cases synthesized dsDNA, were assembled in a Gibson reaction with *pHD-T2A-Gal4-ScarlessDsRed-GMR-eya(shRNA)* that had been digested with EcoRI-HF (NEB, R3101S), or *pHD-T2A-Gal4-ScarlessDsRed-GMR-eya(shRNA)* that had been digested with AscI/SacI-HF (NEB, R0558S, R3156S). In some cases, correctly assembled donor plasmids were screened by colony PCR using common primer pairs that flank the homology arms.

#### sgRNA plasmid cloning

sgRNA target sites were selected using the Find CRISPRs tool (https://www.flyrnai.org/crispr3/web/)^96^. Plasmids encoding sgRNAs were constructed by annealing oligos and ligating into *pCFD3* (Addgene # 49410)^97^ digested with BpiI (Thermo Fisher, ER1011), or PCR amplifying dual sgRNA sequences and Gibson cloning into *pCFD5* (Addgene # 73914)^98^ digested with BpiI.

### S2R+ cell culture

*Drosophila* S2R+ cells^99^ were cultured at 25 °C using Schneider’s media (21720-024; Thermo Fisher Scientific) with 10% fetal bovine serum (A3912; Sigma) and 50 U/mL penicillin-streptomycin (15070-063; Thermo Fisher Scientific). Cells were transfected using Effectene (301427; Qiagen) following the manufacturer’s instructions.

For testing TurboID constructs by imaging, S2R+ cells were transfected with *pMT-sfGFP-TurboID* or *pMT-sfGFP-TurboID-ER* plasmids in 6-well plates. Four days after transfection, 100uM CuSO4 and 50uM biotin (Sigma, B4639-1G) was added to induce gene expression and promote biotinylation, respectively. 24 hours later, cells were resuspended, diluted to 1x10^6 cells/ml, and transferred to 384 well glass plates (PerkinElmer, 6007558).

For testing labeling of Spz and hhN secreted proteins, S2R+ cells were transfected in 6-well plates with *pAct-Gal4* (Y. Hiromi, National Institute of Genetics, Mishima, Japan), and *pUAS-spatzle-HA* or *pUAS-hhN-HA*, and *pMT-GFP-V5* or *pMT-sfGFP-TurboID-ER*.

*S2R+-MT-sfGFP-TurboID* (DGRC #350) and *S2R+-MT-sfGFP-TurboID-ER* (DGRC #351) stable cell lines were generated by transfecting S2R+ with *pMT-sfGFP-TurboID* or *pMT-sfGFP-TurboID-ER* plasmids in 6-well plates. These *pMK33*^52^-derived plasmids contain a Hygromycin resistance gene and a *Metallothionein* promoter to induce gene expression. After 4 days, transfected cells were selected with 200 µg/ml Hygromycin (Calbiochem; 400051-1MU) in Schneider’s medium for approximately 1 month. *S2R+-MT-Myc-BirA*G3* and *S2R+-MT-Myc-BirA*G3-ER* cell lines were isolated in the same manner.

For western analysis of biotinylated proteins, *S2R+-MT-sfGFP-TurboID* or *S2R+-MT-sfGFP-TurboID-ER* cells were grown in T25 flasks until near confluency (8x10^6 cells/ml). 100uM CuSO4 and 50uM biotin (Sigma, B4639-1G) was added to induce gene expression and promote biotinylation, respectively. 24 hours later, cells and culture media were collected separately. First, 1ml of culture media was collected and centrifuged at 1000g for 5 min to pellet remaining cells. The cell-free media was filtered through a 0.2µm centrifuge filter (VWR, 525-0017) and concentrated with a 0.5ml 3 kDa centrifuge filter (Sigma, UFC500324) at 14,000g. To help remove excess biotin from the sample, 500µL of PBS was added to the retentate on the column and the sample was centrifuged again. This biotin wash was performed 2x. The retentate was recovered (∼25µL) and protein concentration determined by BCA assay (Thermo Fisher, 23227). Second, the transfected cells were resuspended, centrifuged for 1000g for 5min, and the supernatant was discarded. The cell pellet was resuspended in PBS and centrifugation repeated, the supernatant discarded, the cell pellet was lysed in RIPA buffer with protease inhibitors (Thermo Fisher, 89901, 87786), and the protein concentration determined by BCA assay.

For small-scale streptavidin pulldown experiments involving transfection (Figure 1E), four days after transfection in 6-well plates, 100uM CuSO4 and 50uM biotin (Sigma, B4639-1G) was added to induce gene expression and promote biotinylation, respectively. For small-scale streptavidin pulldown experiments involving stable cell lines (**Supplemental** Figure 1D), cells were grown to ∼8x10^6 cells/ml in 6-well plates and 100uM CuSO4 and 50uM biotin was added to induce gene expression and promote biotinylation, respectively. 24 hours later, 1ml of culture media was collected and centrifuged at 1000g for 5 min to pellet remaining cells. The cell-free media was filtered through a 0.2µm centrifuge filter (VWR, 525-0017) and concentrated with 0.5ml 3 kDa centrifuge filter (Sigma, UFC500324) at 14,000g. To help remove excess biotin from the sample, 500µL of PBS was added to the retentate on the column and the sample was centrifuged again. This biotin wash was performed 2x. The retentate was recovered (∼25µL) and protein concentration determined by BCA assay (Thermo Fisher, 23227).

For large-scale streptavidin pulldown experiments (**Supplemental** Figure 1C), *S2R+-MT-sfGFP-TurboID* and *S2R+-MT-sfGFP-TurboID-ER* cells were grown in 182cm^2^ cell culture flasks (hold 30ml volume) until nearly confluent (8x10^6 cells/ml). 100uM CuSO4 and 50uM biotin (Sigma, B4639-1G) was added to induce gene expression and promote biotinylation, respectively. 24 hours later, 30ml of culture media was collected and centrifuged at 1000g for 5 min to pellet remaining cells. The cell-free media was filtered through a 0.2µm centrifuge filter (Thomas Scientific, 229710) and concentrated with 15ml 3 kDa centrifuge filter (Sigma, UFC900324) at 4,000g. To help remove excess biotin from the sample, 15ml of PBS was added to the retentate on the column and the sample was centrifuged again. This biotin wash was performed 2x. The retentate was recovered (200-500µL) and protein concentration determined by BCA assay (Thermo Fisher, 23227).

### Antibody staining and imaging

S2R+ cells in 384 well plates were fixed in 4% paraformaldehyde (Electron Microscopy Sciences, 15710) for 30min, washed with PBS with 0.1% Triton X-100 (PBT) three times for 5 min each, stained with primary antibodies overnight at 4°C, washed with PBT, stained with secondary antibodies for 2hr at 25°C, and washed with PBS. Plates were imaged on an IN Cell Analyzer 6000 (GE Healthcare) using a 60x objective. Images were processed using Fiji software.

Antibodies used:

mouse anti-Cnx99A (1:10, DSHB, Cnx99A 6-2-1)

chicken anti-GFP (1:1000, AVES labs GFP1020)

streptavidin-647 (1:500, Thermo Fisher, S21374)

goat anti-chicken 488 (1:500, Thermo Fisher, A-11039)

donkey anti-mouse 488 (1:500, Molecular Probes A-21202)

rabbit anti-myc (1:250, Cell Signaling 71D10)

donkey anti-rabbit 488 (1:500, Molecular Probes A21206)

Whole larvae fluorescence was imaged by immobilizing 3^rd^ instar larvae in a drop of PBS in a glass-well dish on a slide warmer (set to maximum) for 5min. Larvae were imaged on a Zeiss Axio Zoom V16 fluorescence microscope. Tissue-Gal4, UAS-GFP-TurboID-ER lines were crossed with *UAS-6x-GFP* to boost fluorescence signal.

For imaging larval tissues, wandering 3rd instar larvae were dissected in PBS and tissues were fixed in 4% paraformaldehyde for 20 min. Tissues were permeabilized in PBT, blocked for 1 hr in 5% normal goat serum (S-1000, Vector Labs) at room temperature, and incubated with anti-GFP antibody overnight at 4°C, washed with PBT, incubated with anti-chicken 488 and phalloidin and DAPI for 4 hr at room temperature, washed with PBT and PBS, and incubated in mounting media (90% glycerol + 10% PBS) overnight at 4 °C. Stained tissues were mounted on a glass slide under a coverslip using vectashield (H-1000, Vector Laboratories Inc). Images of stained tissues were acquired on a Zeiss 780 confocal microscope. Images were processed using Fiji software.

Antibodies/stains used:

chicken anti-GFP (1:1000, AVES labs GFP1020)

goat anti-chicken 488 (1:500, Thermo Fisher, A-11039)

phalloidin-660 (1:40, Invitrogen, A22285)

DAPI (1:1000, Invitrogen, D1306)

### Western blotting and SDS-PAGE gel staining

Protein or cell samples were denatured in 2x SDS Sample buffer (100 mM Tris-CL pH 6.8, 4% SDS, 0.2% bromophenol blue, 20% glycerol, 0.58 M β-mercaptoethanol) by boiling for 10 min. Denatured proteins and Pageruler Prestained Protein Ladder (Thermo Fisher Scientific 26616) were loaded into 4–20% Mini-PROTEAN TGX gels (Biorad 4561096) using running buffer (25 mM Tris, 192 mM glycine, 0.1% SDS, pH 8.3). Equal amounts of denatured proteins were loaded per lane unless otherwise noted. Gels were run at 100–200 V in a Mini-PROTEAN Tetra Vertical Electrophoresis Cell (Biorad 1658004).

Denatured proteins loaded per lane:

Figure 1C-D – 15µg Figure 2D – 3.6µg

Supplemental Figure 1B –15µg

Supplemental Figure 1C - Not determined due to sample buffer elution. 20µL loaded.

Supplemental Figure 1D – Input: 5.4µg

Supplemental Figure 1D – Flowthrough: same volume as input

Supplemental Figure 1D – Pulldown: Not determined due to sample buffer elution. 20µL loaded.

Supplemental Figure 2A – Dilution series, µg loaded indicated on figure.

Supplemental Figure 2B - Not determined due to sample buffer elution. 20µL loaded. Supplemental Figure 2C – 20µg

Supplemental Figure 3A – Not determined due to sample buffer elution. 20µL loaded.

Supplemental Figure 3B – Not determined due to sample buffer elution. 20µL loaded. Figure 5E – 3.6µg

Supplemental Figure 11 – 3.6µg

For large-scale pulldown/M.S. involving excised Coomassie-stained gel bands, 5/6^th^ of the elution was ran on a gel and individual protein bands were excised, whereas 1/6^th^ of the elution was ran 1cm into the gel the entire lane excised.

For western blotting, proteins were transferred to Immobilon-FL PVDF (Millipore IPFL00010) in transfer buffer (25 mM Tris, 192 mM glycine) using a Trans-Blot Turbo Transfer System (Biorad 1704150) (Standard SD program). When applicable, blots were stained with Ponceau S Solution (Sigma, P7170) for 5min, washed with ddH20, imaged on a ChemiDoc MP Imaging System (BioRad), and destained in 0.1M NaOH. Blots were incubated in TBST (1 x TBS + 0.1% Tween20) for 20 min on an orbital shaker, blocked in 3% BSA (Sigma, A3912) in TBST for 1 hour at room temperature, and incubated with primary antibody diluted in blocking solution overnight at 4 °C. Blots were washed with TBST and incubated in secondary antibody in blocking solution for 4 hour at room temperature. Blots were washed in TBST before detection of proteins. HRP-conjugated streptavidin or secondary antibodies were visualized using ECL (34580, ThermoFisher). Fluorophore-conjugated secondary antibodies were visualized using fluorescence. Blots were imaged on a ChemiDoc MP Imaging System (BioRad). Target proteins were reprobed by incubating blots with stripping buffer (Thermo Scientific 46430) following the manufacturer’s instructions, re-blocked in 3% BSA in TBST, and incubated with primary antibody overnight as described.

Antibodies used:

Streptavidin-HRP (0.3µg/ml, Thermo Fisher, S911)

Rhodamine Anti-Actin (1:10,000, Biorad 12004163)

rabbit anti-GFP (1:5,000, Invitrogen a6455)

goat anti-rabbit 800 (1:5,000, Thermo Fisher, A32730)

chicken anti-HA (1:5,000, Aves, ET-HA100)

goat anti-chicken-HRP (1:1,000, Sigma, SAB3700199)

rabbit anti-LSP-1γ (1:5000) ^47^

rabbit anti-myc (1:2000, Cell Signaling 71D10)

donkey anti-rabbit-HRP (1:3000, Amersham NA934)

SDS-PAGE gels were stained using silver staining or coomassie staining. For silver-stained gels, we used the Pierce Silver Stain Kit (Thermo Fisher, 24612). For coomassie staining, SDS-PAGE gels were fixed in fixing solution (40% Methanol/10% acetic acid) for 1 hour, stained in 0.25% Coomassie Brilliant Blue R-250 in fixing solution for 4 hours, and destained in fixing solution. Stained gels were imaged on a ChemiDoc MP Imaging System. Gel lanes and bands were cut out using a clean razor blade.

### Streptavidin pulldowns

Streptavidin pulldowns were performed as previously^6^ with modifications noted.

For small-scale pulldowns (i.e. for western analysis), 360µg of sample was resuspended in 500µL RIPA buffer + protease inhibitors (Thermo Fisher, 89901, 87786). 7.5µL of resuspended sample was removed as input and boiled with 7.5µL 4x SDS Sample Buffer (200 mM Tris-CL pH 6.8, 8% SDS, 0.4% bromophenol blue, 40% glycerol, 1.16 M β-mercaptoethanol). 30µL of magnetic streptavidin beads (Pierce 88817) were washed in RIPA buffer using a magnetic rack (Bio-Rad, 1614916), combined with the sample, and allowed to bind the beads for 1 hour at room temperature. Note that **Supplemental** Figure 1D used the small-scale pulldown protocol.

For large-scale pulldowns (i.e. for LC-MS/MS), 3mg of sample was diluted with RIPA buffer + 1x protease inhibitors to 500µL total volume. 5µL of this sample input was retained and boiled with 4x sample buffer for later SDS-PAGE analysis. 250µL of magnetic streptavidin beads were washed in RIPA buffer using a magnetic rack, combined with the sample, and allowed to bind the beads for 1 hour at room temperature. Note that **Supplemental** Figure 1C used the large-scale pulldown protocol.

Sample + beads were incubated rocking for one hour at room temperature. Beads were washed sequentially to remove nonspecific binders (1ml for each wash): twice with RIPA buffer, once with 1M KCL, once with 0.1 M Na2CO3, once with 2 M urea in 10 mM Tris-HCl (pH 8.0), and twice with RIPA buffer.

To reduce non-specific binding to beads, we tested alternative urea wash steps, which ranged from 2M, 4M, 6M, or 8M urea and 0%, 1%, or 2% SDS. We used 4M urea + 2% SDS in 10 mM Tris-HCl pH 8.0 for pulldowns for the full 10-tissue secretome map using TMT labeling and LC-MS/MS.

For elution of biotinylated proteins from beads to run on an SDS-PAGE gel, beads were boiled in 30µL of 3x SDS Sample buffer with biotin and DTT (150 mM Tris-CL pH 6.8, 6% SDS, 0.3% bromophenol blue, 30% glycerol, 0.87 M β-mercaptoethanol, 2 mM biotin, 20 mM DTT) for 10 min.

For submission of biotinylated proteins for on-bead digestion, we suspended washed beads in 500µL in PBS (for LC-MS/MS using spectral counting) or RIPA buffer (for LC-MS/MS using TMT labeling).

### Protein localization prediction

Uniprot entry #s from LC-MS/MS data were input into the Uniprot Retrieve/ID mapping tool (https://www.uniprot.org/id-mapping), and exported fields “Subcellular Location”, “Topological domain”, “Transmembrane”, and “Signal”. For the “Subcellular Location” and “Topological domain” fields, secreted and/or transmembrane proteins were identified using keywords “secreted” or “extracellular”. For the “Transmembrane” and “Signal” fields, secreted and/or transmembrane proteins were identified by the presence of a predicted transmembrane domain or signal sequence, respectively.

Uniprot entry #s were converted to FlyBase gene names (FBgns), input into the Flybase Batch Download tool (http://flybase.org/batchdownload), and exported the field “Gene Ontology(GO): Cellular Component”. Secreted and/or transmembrane proteins were identified using keywords “extracellular space ; GO:0005615” or “extracellular region ; GO:0005576”.

FBgns were compared with predictions of “Secreted proteins” and “Trans-membrane proteins” the Gene List Annotation for Drosophila (GLAD) tool (https://www.flyrnai.org/tools/glad/web/)^100^.

FBgns were converted to Flybase polypeptides (FBpp) for all protein isoforms (http://flybase.org/download/sequence/batch) and used as input for SignalP 6.0^101^, SignalP 5.0^102^,TMHMM 2.0^103^, DeepTMHMM^104^, and DeepLoc 2.0^105^. In all cases, the default threshold of each software package was used to classify whether a protein had a secretion signal sequence (SignalP), transmembrane domain (TMHMM and DeepTMHMM), or “Extracellular” localization (DeepLoc). To maximize identifying domains suggestive of secretion, any single protein isoform that scored as secreted, transmembrane, or extracellular was used as representative of the gene encoding all protein isoforms.

We classified proteins as either “secreted” or “transmembrane” by comparing localization predictions from Uniprot, Flybase, GLAD, and bioinformatic prediction software. For example, proteins that contained a secretion signal sequence with no transmembrane domain were classified as “secreted”, and proteins that contained a transmembrane domain as “transmembrane”.

### Homology searching

Fly-human homologs were identified using the DIOPT Ortholog Finder (https://www.flyrnai.org/diopt) version 9.1^106^. Homologs were defined as having a DIOPT score of 3 or higher.

### Drosophila genetics

Flies were maintained on standard fly food at 25 °C.

Fly stocks were obtained from the Perrimon lab collection, stock centers, donating labs, or generated in this study (see below).

Perrimon Lab stocks:

*Lpp-Gal4* (Chr. III)

*Hml-Gal4* (Chr. X)

*yw; nos-Cas9 attp40* (Chr. II)

*yw; nos-Cas9 attp2* (Chr. III)

Bloomington Drosophila Stock Center:

*w*^1118^ (5905)

*repo-Gal4* (7415)

*promE-Gal4* (65405)

*elav-Gal4* (8760)

*MS10906-Gal4* (8860)

*mef2-Gal4* (27390)

w[1118]; Herm[3xP3-ECFP,alphatub-piggyBacK10]M6; MKRS/TM6B, Tb[1]) (32070) yv,P[y[+t7.7]=nos-phiC31\int.NLS]X; P[y[+t7.7]=CaryP]attP40 (25709) P[y[+t7.7]=nos-phiC31\int.NLS]X, y(1) sc(1) v(1) sev(21); P[y[+t7.7]=CaryP]attP2 (25710)

UAS-6x-GFP (52262)

Kyoto Stock Center:

btl-Gal4 (105276)

myo1a-Gal4 (112001)

Other sources:

*4xHand-Gal4107*

This study (BDSC stock # indicated):

*w; UAS-sfGFP-TurboID-ER Chr.II* (BDSC 606643)

*w;; UAS-sfGFP-TurboID-ER Chr.III* (BDSC 606644)

*w;; Mef2-Gal4, UAS-sfGFP-TurboID-ER/TM3, Sb (BDSC 606645)*

*w, MS1096-Gal4; UAS-sfGFP-TurboID-ER (BDSC 606646)*

*w;; 4xHand-Gal4, UAS-sfGFP-TurboID-ER/CyO (BDSC 606647)*

*w, hml-Gal4; UAS-sfGFP-TurboID-ER (BDSC 606648)*

*w;; promE-Gal4, UAS-sfGFP-TurboID-ER/TM3 (BDSC 606649)*

*w; Myola-Gal4, UAS-sfGFP-TurboID-ER/CyO (BDSC 606650)*

*w;; lpp-Gal4, UAS-sfGFP-TurboID-ER/TM3 (BDSC 606651)*

*w;; btl-Gal4, UAS-sfGFP-TurboID-ER/TM3 (BDSC 606652)*

*w;; elav-Gal4, UAS-sfGFP-TurboID-ER/TM3 (BDSC 606653)*

*w;; repo-Gal4, UAS-sfGFP-TurboID-ER/TM3 (BDSC 606695)*

*yw;; UAS-CG6867-sfGFP (BDSC 606654)*

See **Supplemental Table 6** for information on knock-in fly strains

Transgenic flies were made by injecting *attB*-containing plasmid at 200 ng/µL into *nos-PhiC31* integrase-expressing embryos that contain an *attP* landing site (*attP40* or *attP2*). The plasmid contains a *white+* marker gene for positive selection. Injected adults were outcrossed to *white-* balancer chromosome lines to isolate transgenic founder flies with *white+* (red) eyes and generate balanced stocks.

Fly strains stably carrying both *Gal4* and *UAS-sfGFP-TurboID-ER* transgenes were generated by genetic crosses between *Gal4* strains with *UAS-sfGFP-TurboID-ER*, using recombination in cases where both transgenes were on the same chromosome. Isolating flies carrying both transgenes was aided by manually selecting larvae or adults that contain GFP fluorescence, using a Zeiss Axio Zoom V16 fluorescence microscope.

Knock-in flies were made by co-injecting sgRNA plasmid and donor plasmid (500 ng/µL total) into *nos-Cas9* embryos. We used a 5:1 ratio of donor:sgRNA plasmid. *nos-Cas9 attp40* was used for targeting genes on Chr. III and *nos-Cas9 attp2* was used for targeting genes on Chr. X, II, and IV. Injected embryos were raised to adult flies, which were outcrossed to *yw* and adult progeny were screened for RFP+ fluorescent eyes on a Zeiss Axio Zoom V16 fluorescence microscope. Injected males were set up as single male crosses to *yw* females, and injected females were set up as pooled crosses with *yw* males. We recorded the number of injected male crosses or pooled female crosses that gave any RFP+ progeny. We also recorded the number of injected male crosses or pooled female crosses that gave any RFP+ progeny that had small eyes. Candidate knock-in flies (RFP+, normal size eyes) were outcrossed to a balancer chromosome to establish a stock, while also collecting RFP+ flies for PCR genotyping. Genomic DNA was isolated from 25 adult RFP+ flies using the Quick-DNA miniprep kit (Zymo, D3024). Candidate RFP+ lines were genotyped by PCR by amplifying and sequencing the region flanking both homology arms. Primers that amplify the homology arm region were designed such that one primer binds to genomic sequence distal to the homology arm and the other primer binds to insert sequence (**Supplemental Figure 10C**). This strategy ensures that the target locus is amplified. Amplified homology arm regions were sanger sequenced to confirm the correct knock-in event. The overall knock-in success rate was calculated as the number of targeting constructs (i.e. unique injection mixes) vs the number of successful knock-in fly strains (**Supplemental Table 6**)

To generate scarless sfGFP-tagged knock-in lines, the *3xP3-RFP* marker was removed by crossing pre-excision alleles to a PBac transposase strain (*Herm[3xP3-ECFP,alphatub-piggyBacK10]M6*), outcrossing F1 progeny to a balancer strain, and selecting F2 progeny that lack the RFP+ eye marker. Candidate sfGFP-tagged alleles were genotyped like as described above, using one primer that binds *GFP* sequence and one that binds distal to the homology arm (**Supplemental Figure 10C**).

### Larval hemolymph isolation

#### Large-scale collection (for streptavidin pulldowns)

To synchronize developmental timing and prevent overcrowding of larvae, we allowed adult flies to lay eggs on food with yeast pellets for 6-8 hours, and progeny were raised at 25°C. Four days later, larvae were floated with 20% Glycerol (Sigma, G7757-1GA) and ∼200 were transferred to fresh fly food containing excess biotin (Sigma, B4639-1G). Biotin fly food was prepared by mixing a biotin stock solution (1mM Biotin in H20) 1:10 with microwaved liquid fly food (final 100µM Biotin). Transferred larvae were raised on biotin fly food at 25°C for 24 hours. Five-day old larvae were floated with 20% Glycerol and washed using a mesh basket (Genesee Scientific, 46-102, 57-107, 57-101) to remove excess biotin fly food stuck to larvae. 1g washed larvae (∼400 larvae) were transferred to a glass dish (Carolina Biological Supply 742300) on ice. Larvae were washed with ice cold PBS (Gibco, 10010023) in the glass dish until no food particles remained, after which larvae were resuspended in 1000µL ice cold PBS.

Larvae were bled using micro-scissors (World Precision Instruments, 501778) to nick posterior cuticle. Larval bleeding for each sample (∼400 larvae) was performed within ∼30min while keeping the glass dish on ice. Larval blood in PBS was filtered through a 100µm cell strainer (Falcon, 352360) to remove carcasses, using ice-cold PBS to gently rinse the glass dish and carcasses in the cell strainer, finishing with no more than 15ml diluted crude larval blood in PBS per sample (∼400 larvae). Crude diluted blood was centrifuged at 250g for 10min, the supernatant was passed through a 0.8µm/0.2µm filter (Pall Corporation 4187) using a 30ml syringe (BD 309650), and the flowthrough centrifuged through a 15ml 3 kDa centrifugal filter unit (Millipore UFC9003) at 4000g. The resulting ∼250µL purified concentrated blood was removed from the centrifugal filter unit and diluted with 500µL RIPA (Thermo Fisher, 89901) + 1.5x protease inhibitors (Thermo Fisher, 87786). Protein concentration was determined using a BCA assay (Thermo Fisher, 23227). The remaining sample was snap-frozen in liquid nitrogen and stored at -80°C.

#### Small-scale collection (for non-pulldowns, for western blotting)

20 3^rd^ instar larvae were washed in cold PBS and transferred to a paper towel to dry for 1 minute. Larvae were transferred to a 1”x1” square of parafilm. Larvae were bled using micro-scissors (World Precision Instruments, 501778) to nick posterior cuticle, and the carcasses were pushed together to pool the bled hemolymph. 2µL of raw hemolymph was removed and transferred to 10µL of cold PBS. The diluted hemolymph was centrifuged at 1000g for 10min at 4°C to pellet cells and tissue. 10µL of supernatant was transferred to a new tube containing 10µL 4x sample buffer and boiled for 10min.

### LC-MS/MS for pilot TurboID-ER labeling experiments in S2R+ cells and hemolymph

Protein samples were reduced with 55 mM DTT, alkylated with 10 mM iodoacetamide (Sigma-Aldrich), and digested overnight with TPCK modified trypsin/LysC (Promega) at pH = 8.3. Peptides were immunoprecipitated with a pTyr antibody, dried out in a SpeedVac, resuspended in 10 μL of 1% Acetonitrile/98.9% Water, 0.1% Formic acid. 3 uL of the digested protein samples were analyzed in positive ion mode via microcapillary tandem mass spectrometry (LC-MS/MS) using a high resolution hybrid QExactive HF Orbitrap Mass Spectrometer (Thermo Fisher Scientific) via HCD with data-dependent analysis (DDA) with 1 MS1 scan followed by 8 MS2 scans per cycle (Top 8). Peptides were delivered and separated using an EASY-nLC1000 nanoflow UPLC (Thermo Fisher Scientific) at 300 nL/min using self-packed 15 cm length × 75 μm i.d. C18 fritted microcapillary columns. Solvent gradient conditions were 90 minutes from 3% B buffer to 38% B (B buffer: 100% acetonitrile; A buffer: 0.9% acetonitrile/0.1% formic acid/99.0% water). MS/MS spectra were analyzed using Mascot Version 2.7 (Matrix Science) by searching the reversed and concatenated *Drosophila* protein Database: the Drosophila_20180328 database (unknown version, 13767 entries) with a parent ion tolerance of 18 ppm and fragment ion tolerance of 0.05 Da. Carbamidomethylation of cysteine (+57.0293 Da) was specified as a fixed modification and oxidation of Methionine (+15.9949 Da), deamidation of Asparagine/Glutamine (+0.984 Da) as variable modifications. Results were imported into Scaffold Q+S 5.0 software (Proteome Software, Inc.) with a peptide threshold of ∼70%, protein threshold of 95%, resulting in a peptide false discovery rate (FDR) of ∼1%.

For tissue-labeling experiments involving whole-lane in-gel digestion, tissue-enriched proteins were calculated as greater than 3 spectral counts from the tissue and at least 0.25 enrichment (# spectral counts from specific tissue / # spectral counts from all tissues).

### Tissue secretome map: On-bead trypsin digestion of biotinylated proteins

On-bead trypsin digestion was performed as described previously^6^. The biotin-labeled proteins bound to magnetic beads were further washed to remove detergent traces. Magnetic beads were immobilized and washed twice with 200 μL of 50 mM Tris HCl buffer (pH 7.5) followed by two washes with 2 M urea/50 mM Tris (pH 7.5) buffer. A partial trypsin digestion was performed to release proteins from the beads by using 80 μL of 2 M urea/50 mM Tris containing 1 mM DTT and 0.4 μg trypsin (Mass Spectrometry Grade, Promega, catalog #V5280) for 1 h at 25 C. Magnetic beads were immobilized and the supernatant containing partially digested proteins was transferred to a fresh tube. The beads were washed twice with 60 μL of 2 M urea/50 mM Tris buffer (pH 7.5), and the washes were combined with the on-bead digest supernatant. Proteins were reduced with 4 mM DTT for 30 min at 25°C with shaking, followed by alkylation with 10 mM iodoacetamide for 45 min in the dark at 25°C. Proteins were completely digested by adding 0.5 μg of trypsin and incubating overnight at 25°C with shaking. Following the overnight incubation, samples were acidified to 1% formic acid (FA) and desalted using stage tips containing 2x C18 discs (Empore, Fisher Scientific catalog # 13-110-019) as described next. The stage tip column was conditioned with 1x 100 μL methanol, 1x 100 μL 50% acetonitrile (ACN) / 1% FA, and 2x 100 μL 0.1% FA washes. Acidified peptides were bound to the column, washed with 2x 100 μL 0.1% FA, and eluted with 50 μL 50%ACN/0.1%FA. Eluted peptides were dried to using a vacuum concentrator.

### Tissue secretome map: TMT-labeling and fractionation of peptides

Desalted peptides were labeled with 10-plex TMT reagents (ThermoFisher Scientific, catalog #9406) and fractionated as described previously^6^. Dried peptides were reconstituted in 100 μL of 50 mM HEPES, labeled using 0.8 mg of TMT reagent in 41 μL of anhydrous acetonitrile for 1 h at room temperature. The TMT reactions were quenched with 8 μL of 5% hydroxylamine at room temperature for 15 minutes with shaking, and the labeled peptides were desalted on C18 stage tips as described above. Peptides were fractionated by basic reverse phase using styrenedivinylbenzene-reverse phase sulfonate (SDB-RPS, Empore, Fisher Scientific catalog # 13-110-023) material in stage tip columns. Two-disc punches were packed in a tip and equilibrated with 50 μL methanol, 50 μL 50% CAN / 0.1% FA, and 2x 75 μL 0.1% FA washes. Peptides were reconstituted in 0.1% FA and loaded into the column, followed by conditioning with 25 μL of 20 mM ammonium formate (AF). Next, peptides were sequentially eluted into 6x 100 μL fractions with 20mM AF and varying concentrations of ACN: 5%, 10%, 15%, 20%, 30%, and 55%. The six peptide fractions were dried by vacuum centrifugation.

### Tissue secretome map: liquid chromatography and mass spectrometry analysis

Desalted peptides were resuspended in 9 μL of 3% ACN/0.1% FA and analyzed by online nanoflow liquid chromatography tandem mass spectrometry (LC-MS/MS) using a Proxeon Easy-nLC 1200 coupled to a Q Exactive HF-X Hybrid Quadrupole-Orbitrap Mass Spectrometer (ThermoFisher Scientific). Four μL of sample from each fraction were injected in a capillary column (360 x 75 µm, 50 °C) containing an integrated emitter tip packed to a length of approximately 25 cm with ReproSil-Pur C18-AQ 1.9 μm beads (Dr. Maisch GmbH, catalog # r119.aq). Chromatography was performed with a 110 min gradient of solvent A (3% ACN/0.1% FA) and solvent B (90% ACN/0.1% FA). The gradient profile, described as min:% solvent B, was 0:2, 1:6, 85:30, 94:60, 95:90, 100:90, 101:50, 110:50. Ion acquisition was performed in data-dependent MS2 (ddMS2) mode with the following relevant parameters: MS1 acquisition (60,000 resolution, 3E6 AGC target, 10ms max injection time) and MS2 acquisition (Loop count = 20, 0.7m/z isolation window, 31 NCE, 45,000 resolution, 5E4 AGC target, 105 ms max injection time, 1E4 intensity threshold, 15s dynamic exclusion, and charge exclusion for unassigned, 1 and >6).

### Tissue secretome map: proteomic data analysis

Collected RAW LC-MS/MS data were analyzed using Spectrum Mill software package v6.1 pre-release (Agilent Technologies). MS2 spectra were extracted from RAW files and merged if originating from the same precursor, or within a retention time window of +/- 60 s and m/z range of +/- 1.4, followed by filtering for precursor mass range of 750-6000 Da and sequence tag length > 0. MS/MS search was performed against a custom concatenated FASTA database containing (1) the *Drosophila melanogaster* UniProt (www.uniprot.org) protein database (UP000000803) downloaded on September 2018, (2) a list of common non-human and non-fly contaminants, and (3) the GFP-TurboID-KDEL protein sequence (**Supplemental File 1**). Search parameters were set to “Trypsin allow P”, <5 missed cleavages, fixed modifications (cysteine carbamidomethylation and TMT10 on N-term and internal lysine), and variable modifications (oxidized methionine, acetylation of the protein N-terminus, pyroglutamic acid on N-term Q, pyro carbamidomethyl on N-term C, and NHS-Biotin on K). Matching criteria included a 30% minimum matched peak intensity and a precursor and product mass tolerance of +/- 20 ppm. Peptide-level matches were validated if found to be below the 1.0% false discovery rate (FDR) threshold and within a precursor charge range of 2-5. TMT reporter ion intensities were corrected for isotopic impurities in the Spectrum Mill protein/peptide summary module using the afRICA correction method which implements determinant calculations according to Cramer’s Rule^108^ and correction factors obtained from the reagent manufacturer’s certificate of analysis (Thermo Fisher, cat. # 90406, lot # UA280170).

A protein-centric summary containing TMT channel intensities was exported from Spectrum Mill for downstream analysis in the R environment for statistical computing. Proteins were filtered to remove those that do not originate from *Drosophila* or the TurboID protein. TMT quantitative values were removed if less than 2 peptides were quantified, or the protein score was below 20 for the plex to avoid low-quality quantification. In addition, proteins with a PaxDB score^109^ greater than 1500ppm to remove extremely abundant contaminant proteins (e.g., ribosomes). Sample agreement and outliers were inspected by principal component analysis. Those with large distances in PC1 and PC2 to the average of the experimental group were flagged and manually inspected. Two outlier samples were detected and removed for downstream analysis. TMT ratios were calculated using the median of all control samples, followed by log2 transformation. Two-sample moderated T-tests were performed using the limma package in R with correction for multiple testing by calculating local FDR^110^, producing protein log_2_ fold-changes (logFC) and q-values for contrasts between samples expressing TurboID and controls. Proteins with missing values in all negative controls within a plex were discarded from logFC calculations. Secretion signal sequences were determined by the Uniprot (www.uniprot.org) “Signal” field annotation. Correlation analysis were performed using Pearson’s correlation by comparing TMT ratios between all samples in a plex (**Supplemental** Figure 5). Pearson’s correlation was also performed between the TMT logFC from the tissue secretome map and the log10 spectral counts from the label-free blood proteome dataset (See Methods below and **Supplemental** Figure 6).

Raw proteomic data will be deposited in PRIDE for public availability upon acceptance of the manuscript and will be made available to reviewers upon request.

### Tissue secretome map: Defining high-confidence tissue secreted factors

High-confidence proteins were defined by using an empirical false-discovery rate criteria using a predefined list of secreted positive control (PC) proteins and non-secreted negative controls (NC) (**Supplemental Table 3A**). The positive control list was obtained from^18^, which contains the following: 1) fly receptors and secreted factors, and 2) fly orthologs of human receptors, secreted proteins, and blood plasma proteins. The negative control list contains proteins from the Uniprot database with “cytoplasm”, “nucleus”, or “mitochondria” subcellular location annotations, and proteins from the GLAD database (https://fgr.hms.harvard.edu/glad)^100^ with mitochondrial or transcription factor/DNA binding annotation. Proteins were sorted in descending order using the logFC and the empirical false discovery rate (FDR = FP / (FP+TP)) was calculated at each logFC threshold. False positives (FP) were defined as proteins in the NC list with a logFC equal or greater than the threshold. True positives (TP) were defined as proteins in the PC list with a logFC equal or greater than the threshold. The empirical FDR was plotted as function of the logFC and inspected to manually select a point that minimizes logFC while preserving a small FDR. Four redundant protein isoforms were removed, which were selected in a way as to not lose tissue-of-origin information. We removed B7Z0B3 (*CG10359*) from muscle, because B7Z0B2 (*CG10359*) was identified from muscle/heart/wingdisc/neurons. We removed B7YZF1 (*CG42336*) from fat, because A1Z8I8 (*CG42336*) was identified from fat/gut. We removed D3PK81 (*CtsF*) from muscle, because Q9VN93 (*CtsF*) was identified from muscle, is a longer isoform, and has a “reviewed” status at Uniprot. We removed Q8IQU1 (*verm*) from trachea, because E1JI40 (*verm*) was identified from trachea/gut/heart/wingdisc.

### Blood proteome: Label-free proteomic analysis of whole blood

Whole blood from *Drosophila* larvae (250µL at 30mg/ml) was digested using the S-Trap Midi Spin Column (Protifi, catalog # C02-midi-10) as per manufacturer recommendation. Blood was diluted to 500ul in SDS solubilization buffer (5% SDS, 50mM TEAB pH 7.5), clarified by centrifugation, and acidified using 50uL of 12% phosphoric acid. The S-trap column was conditioned with 3.3mL S-Trap buffer (90% MeOH, 100mM TEAB pH 7.1), samples was loaded by centrifugation, and washed with 3x 3mL S-Trap buffer. Digestion was performed overnight with 1:50 wt:wt trypsin and 1:50 wt:wt LysC in 50mM TEAB. Peptides were eluted sequentially with 500uL each of 50mM TEAB, 0.2% FA, and 0.2%FA/50%ACN. A total of 100µg were lyophilized, resuspended in buffer A (5 mM ammonium formate, pH 10, in 2% acetonitrile), and fractionated by high pH reversed phase separation using a 3.5 um Agilent Zorbax 300 Extend-C18 column (2.1 mm ID x 250 mm length). Samples were loaded onto the column and separated at a flow rate of 1ml/min in a 96 min gradient with the following concentrations of solvent B (5 mM ammonium formate, pH 10.0 in 90% vol/vol MeCN) 16%B at 13min, 40%B at 73min, 44%B at 77min, 60%B at 82min, 60%B at 96min. A total of 96 fractions were collected and concatenated non-sequentially into a final 24 fractions for proteomic analysis. Fractions were dried via vacuum centrifugation.

### Blood proteome: liquid chromatography and mass spectrometry analysis

Dried peptides were reconstituted at an estimated concentration of 0.5 µg/ul in 3% ACN/0.1% FA and analyzed by online nanoflow liquid chromatography tandem mass spectrometry (LC-MS/MS) using a Proxeon Easy-nLC 1200 coupled to a Q Exactive HF-X Hybrid Quadrupole-Orbitrap Mass Spectrometer (ThermoFisher Scientific). One μg of sample from each fraction was in a column setting similar to the one above. Chromatography was performed with a 110 min gradient of solvent A (3% ACN/0.1% FA) and solvent B (90% ACN/0.1% FA). The gradient profile, described as min:% solvent B, was 0:2, 1:6, 85:30, 94:60, 95:90, 100:90, 101:50, 110:50. Ion acquisition was performed in data-dependent MS2 (ddMS2) mode with the following relevant parameters: MS1 acquisition (60,000 resolution, 3E6 AGC target, 10ms max injection time) and MS2 acquisition (Loop count = 20, 0.7m/z isolation window, 28 NCE, 15,000 resolution, 1E5 AGC target, 50 ms max injection time, 1E4 intensity threshold, 7s dynamic exclusion, and charge exclusion for unassigned, 1 and >6).

### Blood proteome: proteomic data analysis

Collected RAW LC-MS/MS data were analyzed using Spectrum Mill software package v6.1 pre-release (Agilent Technologies). MS2 spectra were extracted from RAW files and merged if originating from the same precursor, or within a retention time window of +/- 60 s and m/z range of +/- 1.4, followed by filtering for precursor mass range of 750-6000 Da and sequence tag length > 0. MS/MS search was performed against a custom concatenated FASTA database containing (1) the *Drosophila melanogaster* UniProt protein database (UP000000803) downloaded on September 2018, (2) a list of common non-human and non-fly contaminants. Search parameters were set to “Trypsin allow P”, <5 missed cleavages, fixed modifications (cysteine carbamidomethylation), and variable modifications (oxidized methionine, acetylation of the protein N-terminus, pyroglutamic acid on N-term Q, pyro carbamidomethyl on N-term C, and NHS-Biotin on K). Matching criteria included a 30% minimum matched peak intensity and a precursor and product mass tolerance of +/- 20 ppm. Peptide-level matches were validated if found to be below the 1.0% false discovery rate (FDR) threshold and within a precursor charge range of 2-5. Non-*Drosophila* proteins and proteins with a single unique peptide were removed from the results. The number of peptide-spectrum matches per protein (spectral counts) were used for semi-quantitative analysis and comparison to enrichments from the TMT-based tissue secretome map proteomics results.

### Pathway enrichment analysis and visualization

Pathway enrichment analysis was performed using g:profiler^111^ with the high-confidence secreted proteins (**Supplemental Table 3B**) as input and all proteins identified across all plexes (**Supplemental Table 3A**) as a custom background. The pathway enrichment results for all tissues were imported into Cytoscape v3.8.2 using the EnrichmentMap application for network visualization and cluster analysis with parameters p-value threshold of 1.0, FDR Q-value threshold of 0.05, and Jaccard Overlap Combined value of 0.375^112^.

### Validating protein tissue-of-origin with transcriptome data

The processed *Drosophila* scRNA-seq datasets, such as FCA datasets and larval wing disc datasets, were obtained from DRscDB database^64^ while tissue-specific bulk RNAseq datasets were obtained from DGET database^113^. The tissue/cell type annotation from public datasets was manually matched to the tissue annotation of the secretome data then the genes expressing in each tissue/cell type were compared to the genes identified in the secretome data.

Overlap of our protein tissue secretome and our 3^rd^ instar larval snRNA-seq data was determined by intersecting lists of FlyBase gene names (FBgn) between the two datasets. We represented LC-MS/MS-identified tissue secreted proteins as FBgns (**Supplemental Table 3B**) to enable matching FBgns from transcriptomic data. For validation of expression in a tissue, we counted the number of “Tissue X”-secreted proteins (**Supplemental Table 3B**) that are expressed in any “Tissue X” clusters. We defined expression as 1% or greater average expression OR 10% or greater percent expressed in **Supplemental Table 5F-H**. For validation of tissue-specific expression, we counted the number of “Tissue X”-specific secreted proteins (**Supplemental Table 3B**) that are marker genes in any “Tissue X” clusters. We defined a marker gene as 0.54 or greater avg_log2FC AND greater than 0.05 p_val_adj. To calculate the combined number of tissue-secreted proteins that have a corresponding encoding gene expressed in same tissue cluster(s), we counted the number of unique proteins expressed in at least one tissue (500) (**Supplemental Table 5I**). To calculate the combined number of tissue-specific-secreted proteins that have a corresponding encoding marker gene for the same tissue cluster(s), we counted the total number of cluster marker overlap proteins (105) (**Supplemental Table 5J**).

The 76 high-confidence tissue-specific secreted proteins were determined by counting all tissue-specific secreted proteins that overlapped a marker gene for the same cluster and were predicted secreted or transmembrane (**Supplemental Table 5K**).

### Nuclei isolation from 3^rd^ instar larvae, encapsulation, and sequencing for snRNA-seq

To synchronize developmental timing and prevent overcrowding of larvae, adult flies (*w1118*) were allowed to lay eggs on food supplemented with yeast pellets for 2-4 hours. Progeny were raised at 25°C. Five days later, 16 wandering third instar larvae (8 male, 8 female) that had not everted their spiracles were collected, washed with PBS, snap-frozen in liquid nitrogen, and stored at −80°C.

Nuclei were isolated following a previously described protocol^58,114^. Briefly, 16 frozen larvae were homogenized in a glass dounce containing 1ml of homogenization buffer (250mM sucrose, 10mM Tris pH 8.0, 25mM KCL, 5mM MgCl2, 0.1% Triton-x 100, 0.5% RNaisin Plus [Promega, N2615], 1x protease inhibitor [Promega, G652A], 0.1mM DTT). Nuclei were released by 20 strokes with a loose pestle, followed by 40 strokes with a tight pestle. Lysate containing nuclei was filtered sequentially through a 35µm cell strainer (VWR, 21008-948) and a 40µm cell strainer (Bel-Art, H13680-0040). A crude nuclei sample was isolated by centrifuging filtered lysate for 10min at 1000g at 4°C, discarding the supernatant, and resuspending the pellet in 1000µL 1xPBS/1%BSA by gentle pipetting. The crude nuclei sample was washed by three additional rounds of centrifugation and resuspension in 1xPBS/1%BSA. The washed nuclei sample was filtered through a 40µm cell strainer (Bel-Art, H13680-0040), labeled with 1µM DRAQ7 Dye (Invitrogen D15106), and 500,000 healthy nuclei were isolated by Sony SH800Z Cell Sorter into 1xPBS. FACS sorted nuclei were centrifuged for 10min at 1000g at 4°C and resuspended in 250µL 1xPBS/1%BSA + 1:1000 DAPI (manufacturer). Nuclei concentration was determined by counting DAPI+ nuclei on a hemocytometer, and sample concentration was adjusted to a final concentration of 1,000 nuclei/µL in 1xPBS/1%BSA.

Nuclei were encapsulated and libraries created using the Chromium Next GEM Single Cell 3’ Reagents Kits v3.1, according to the 10X genomics protocol (Chromium Next GEM Single Cell 3’_v3.1_Rev_D). Samples were sequenced on an Illumina NovaSeq 6000 S4. (Following could be the data processing and analysis)

### Analysis of snRNA-seq data

Sequence alignment, cell calling, ambient RNA correction and doublet removal were performed on eight replicates separately and before pooling for batch correction, dimension reductions and clustering. Sequence alignment and cell calling were performed using CellRanger v7.1.0 (10X genomics) in ‘count’ mode using --force-cells=10000. DecontX was performed under default setting and used empty drops from each sample as background. After background correction, cells with less than 50 genes or 80 UMIs were removed. Cells with log10GenesPerUMI>1 and UMI larger than 100,000 were also removed. DoubletFinder with 5% doublet formation rate (according to 10X suggestion) were used to remove potential multiplets from downstream analysis. After these quality control steps, about 30K high quality cells from all samples were retained for downstream analysis. Harmony was performed on the retained cells and top 30 PC of the harmony corrected PCs were used for UMAP visualization and Leiden clustering. Cells were grouped and annotated under clustering resolution=1 for best separation of rarer cell types by top marker genes. The web-based portal for data mining was implemented (https://www.flyrnai.org/scRNA/body_larvae/).

---

**Supplemental Figure 1:**
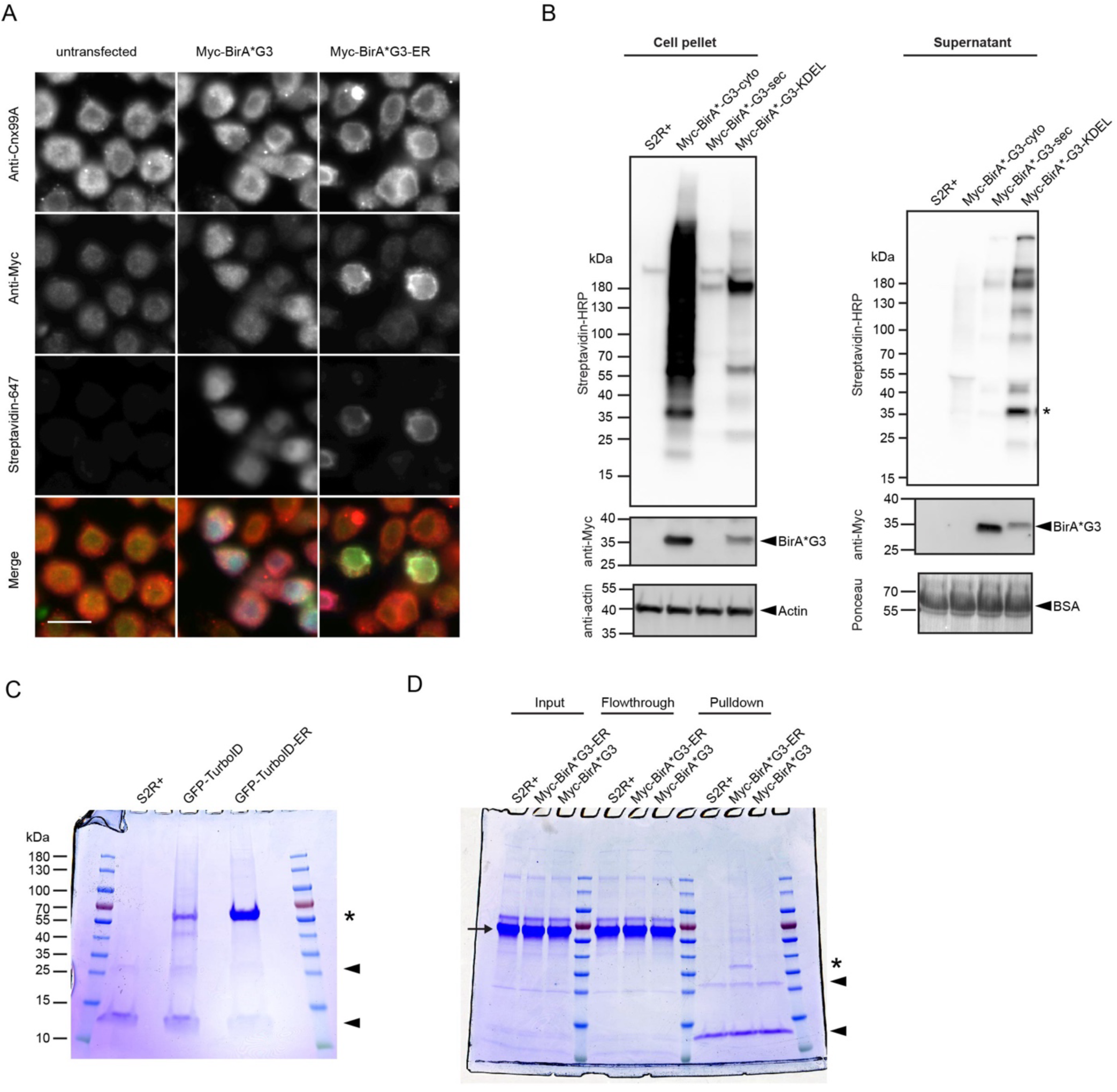
Related to Figure 1. (A) Confocal microscopy of S2R+ cells transfected with *pMT-Myc-BirA*G3* or *pMT-Myc-BirA*G3-ER*. ER detected with Anti-Cnx99A (red), biotinylated proteins detected with streptavidin-647 (blue), and GFP-TurboID detected with anti-GFP (green). Scale bar is 10µm. Red and green channel signal intensity is the same for all samples. Blue channel signal for Myc-BirA*G3 sample was lowered to match the signal intensity of Myc-BirA*G3-ER sample (B) Western blot of cell lysates or media from S2R+ cells transfected with *pMT-Myc-BirA*G3* or *pMT-Myc-BirA*G3-ER*. Each lane loaded 15µg protein. Arrowheads indicate expected band, asterisk indicates auto-biotinylated Myc-BirA*G3. (C) Coomassie-stained SDS-PAGE gel of protein streptavidin-pulldown from 30ml media from *pMT-GFP-TurboID* or *pMT-GFP-TurboID-ER* stable cell lines. Arrowheads indicate streptavidin monomer and dimer. Asterisk indicates auto-biotinylated GFP-TurboID. Note this used the large-scale pulldown in the methods. (D) Coomassie-stained SDS-PAGE gel of protein streptavidin-pulldown from 1ml media from *pMT-GFP-TurboID* or *pMT-GFP-TurboID-ER* stable cell lines. Gel includes input and post-bead flowthrough. Arrowheads indicate streptavidin monomer and dimer. Asterisk indicates auto-biotinylated Myc-BirA*G3. Arrow indicates bovine serum albumin (BSA) from the fetal bovine serum (FBS) in cell culture media. Note this used the small-scale pulldown in the methods.

**Supplemental Figure 2:**
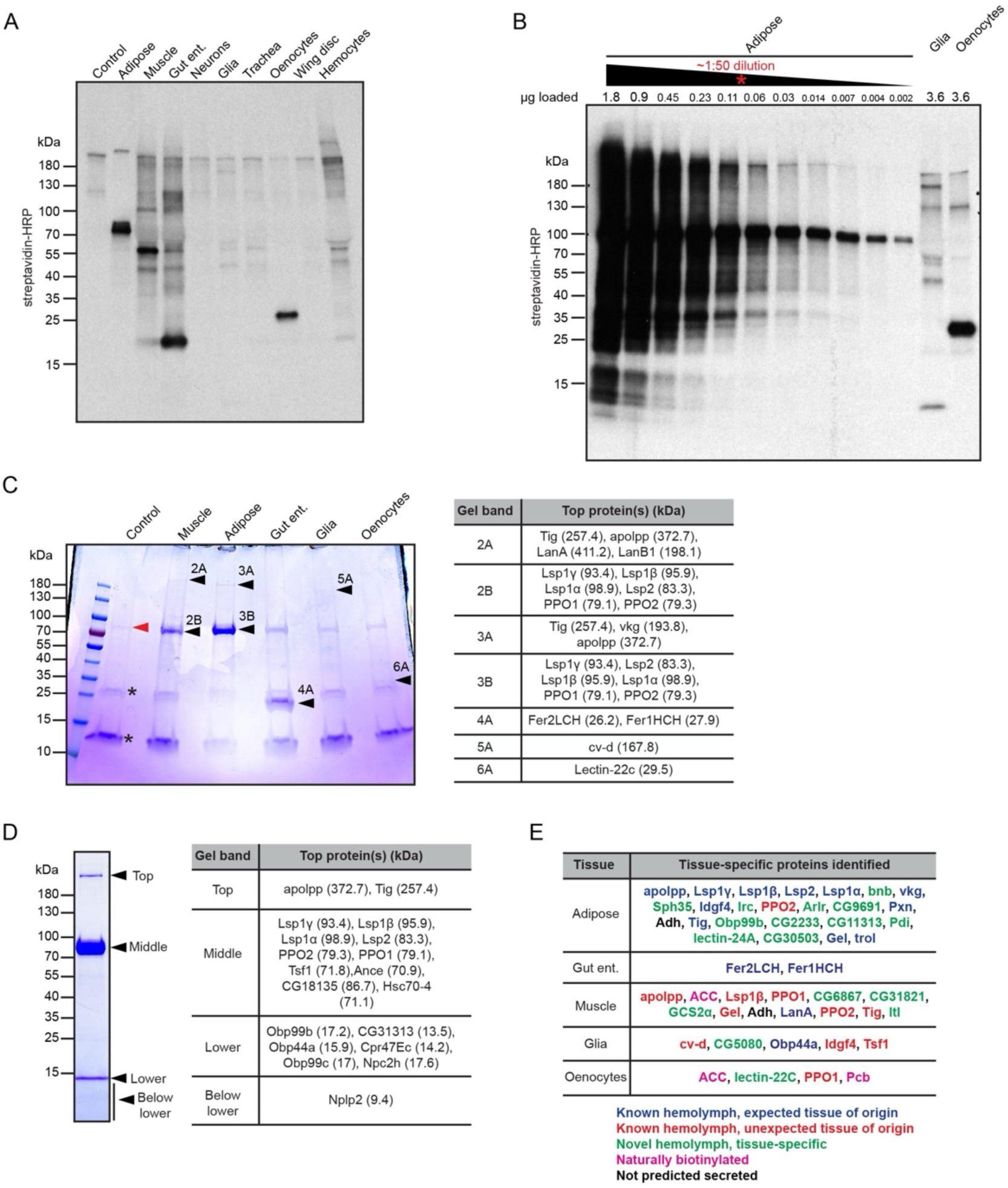
Related to Figure 2. (A) Western blot of biotinylated proteins in purified hemolymph from 3^rd^ instar larvae expressing GFP-TurboID-ER in specific tissues. All lanes loaded 3.6µg protein, except the adipose labeling sample (72ng). Same blot as Figure 2D, lower exposure. (B) Western blot of biotinylated proteins in purified hemolymph from 3^rd^ instar larvae expressing GFP-TurboID-ER in specific tissues. Serial dilution of adipose labeling sample to identify 72ng as comparable signal intensity to glia and oenocyte labeling samples. (C) Coomassie-stained SDS-PAGE gel of protein streptavidin-pulldown from purified hemolymph from 3^rd^ instar larvae expressing GFP-TurboID-ER in specific tissues (left). Black arrowheads indicate protein bands excised for mass spectrometry. Red arrowhead indicates likely non-specific pulldown of LSPs. Asterisks indicate streptavidin monomer and dimer. Table (right) shows top proteins identified. (D) Coomassie-stained SDS-PAGE gel of purified hemolymph from 3^rd^ instar larvae (left). Black arrowheads indicate protein bands excised for mass spectrometry. Table (right) shows top proteins identified. (E) Table of 48 tissue-enriched proteins identified from tissue-labeling streptavidin pulldown and whole-lane gel excision. Enriched proteins defined as ≥0.25 spectral counts/sum spectral counts all experimental samples, and at least 3 spectral counts in enriched tissue. Previous characterization determined using Flybase (”hemolymph” GO term) and literature searching.

**Supplemental Figure 3:**
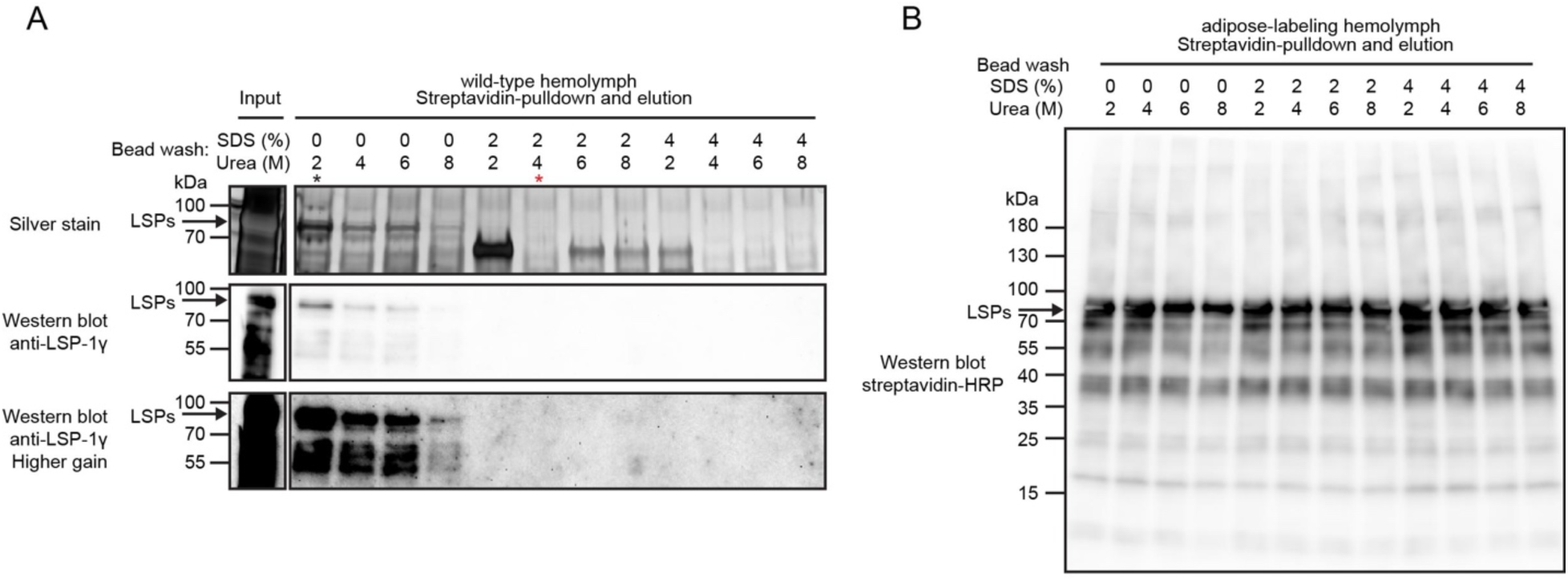
Improved streptavidin-bead washing. (A) Silver stained SDS-PAGE gel (top) and western blot (middle/bottom) of eluted proteins after streptavidin pulldown of purified hemolymph from wild-type 3^rd^ instar larvae. Arrow indicates molecular weight of LSPs. Different beads washing conditions are indicated. Improved washing condition uses 2% SDS and 4M Urea (red asterisk) compared to previous washing condition of 2M Urea (black asterisk). (B) Western blot of eluted biotinylated proteins after streptavidin pulldown of purified hemolymph from 3^rd^ instar larvae with adipose labeling. Arrow indicates molecular weight of LSPs. Different beads washing conditions are indicated.

**Supplemental Figure 4:**
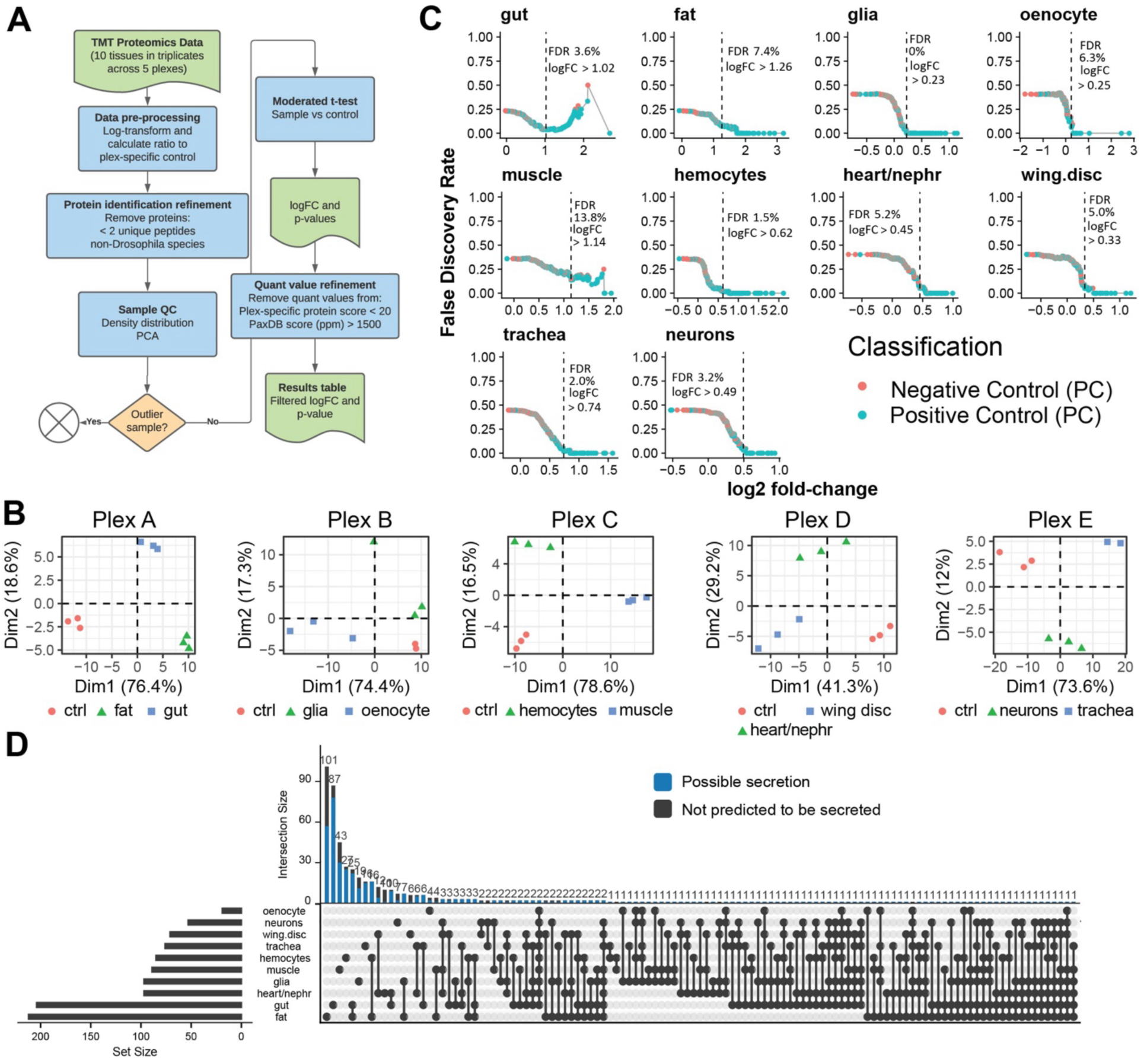
Related to Figure 3. (A) Workflow for data processing and filtering criteria of proteomics data from tissue secreted proteins. (B) Principal component analysis plots for each multiplexed set of samples show cluster of biological replicate samples. (C) Scatter plots showing empirical false discovery rate as a function of log2 fold-change cutoffs using negative and positive controls for classification. The dotted line indicates manually selected point that tries to minimize the fold-change threshold at an acceptable empirical false discovery rate. (D) Upset plot showing the number of proteins identified as secreted by a unique tissue or by multiple tissues. The blue portion of the bar indicates the proportion of proteins annotated as secreted of transmembrane for each intersect.

**Supplemental Figure 5:**
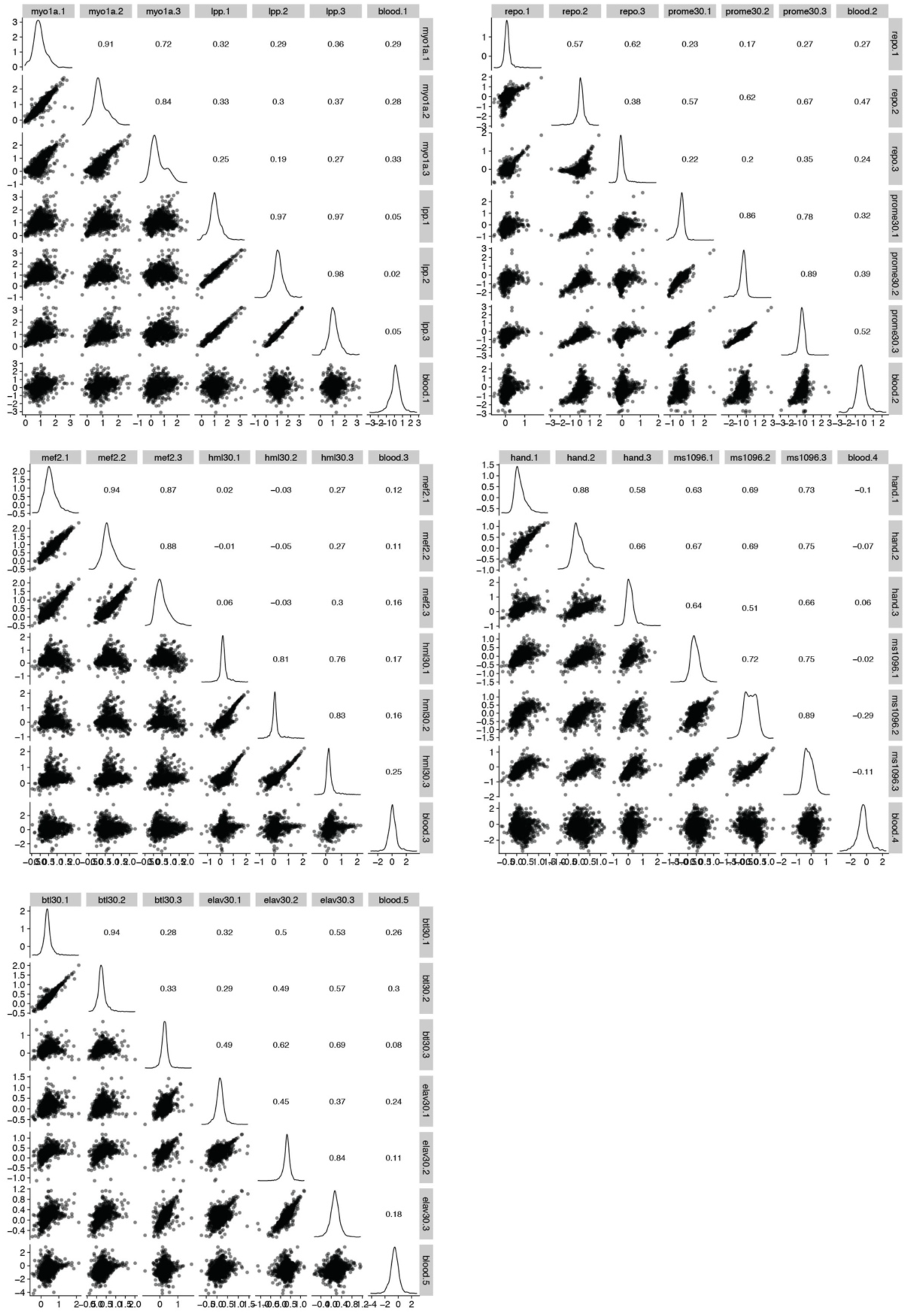
Related to Figure 3. Pairwise plot matrix between samples in the tissue secretome dataset. Pairwise TMT ratio data was plotted within each TMT plex experiment. Scatter plots are shown below the diagonal, density distributions on the diagonal, and Pearson correlation values are shown above the diagonal.

**Supplemental Figure 6:**
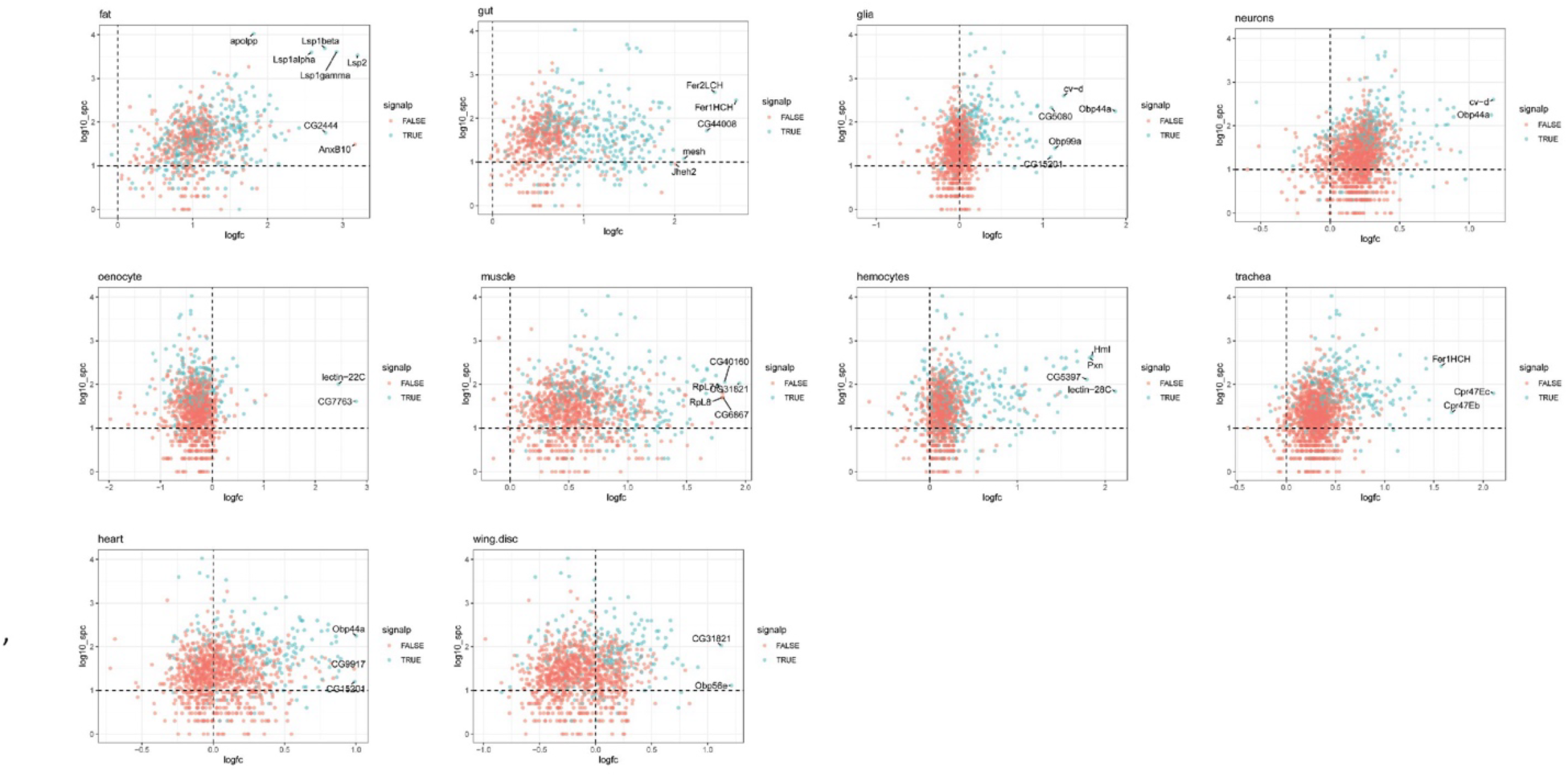
Related to Figure 3. Scatter plot showing log_2_ fold-change from the tissue secretome dataset (tissue/control; logfc) vs the label-free protein abundance from the whole blood proteome dataset as log_10_ spectral counts (log10_spc). Color indicates the content of a signal peptide. Signal peptide annotation by Uniprot. Proteins with a large logfc are highlighted.

**Supplemental Figure 7:**
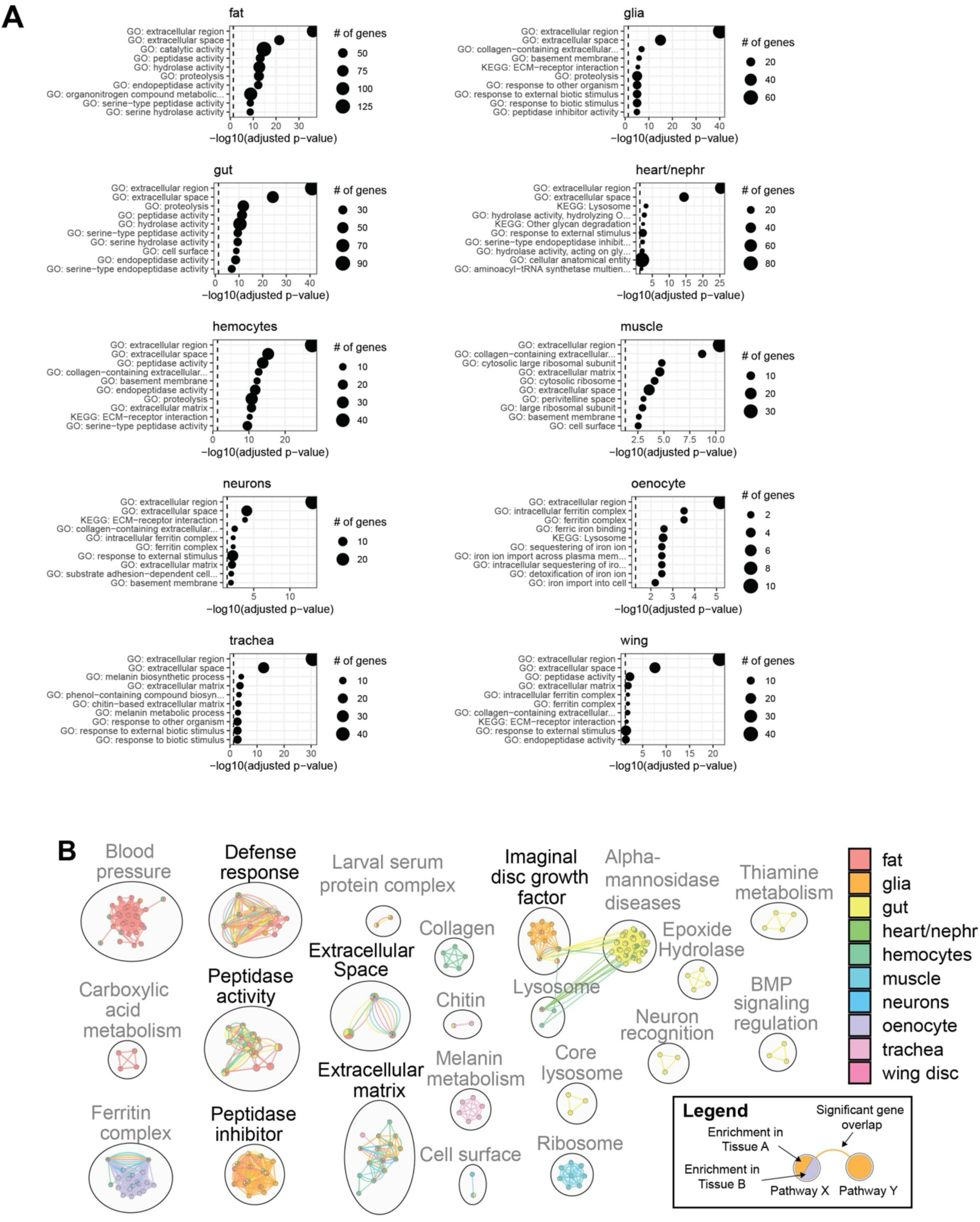
Related to Figure 3. **(A)** Dot plot showing the top 10 pathways from pathway enrichment results for each tissue. Size indicates the number of genes in the pathway. **(B)** Network summary of pathway enrichment results. Nodes indicate pathways with FDR-adjusted p-value below 0.05 colored by the tissue in which the enrichment was observed. Edges indicate that two pathways share a large proportion of genes. Clusters of nodes indicate pathways with similar gene sets with a manual annotation of the common biological theme. Clusters of particular interest are highlighted: immune response, peptidase activity, peptidase inhibitor, extracellular space, extracellular matrix, and imaginal disc growth factor.

**Supplemental Figure 8:**
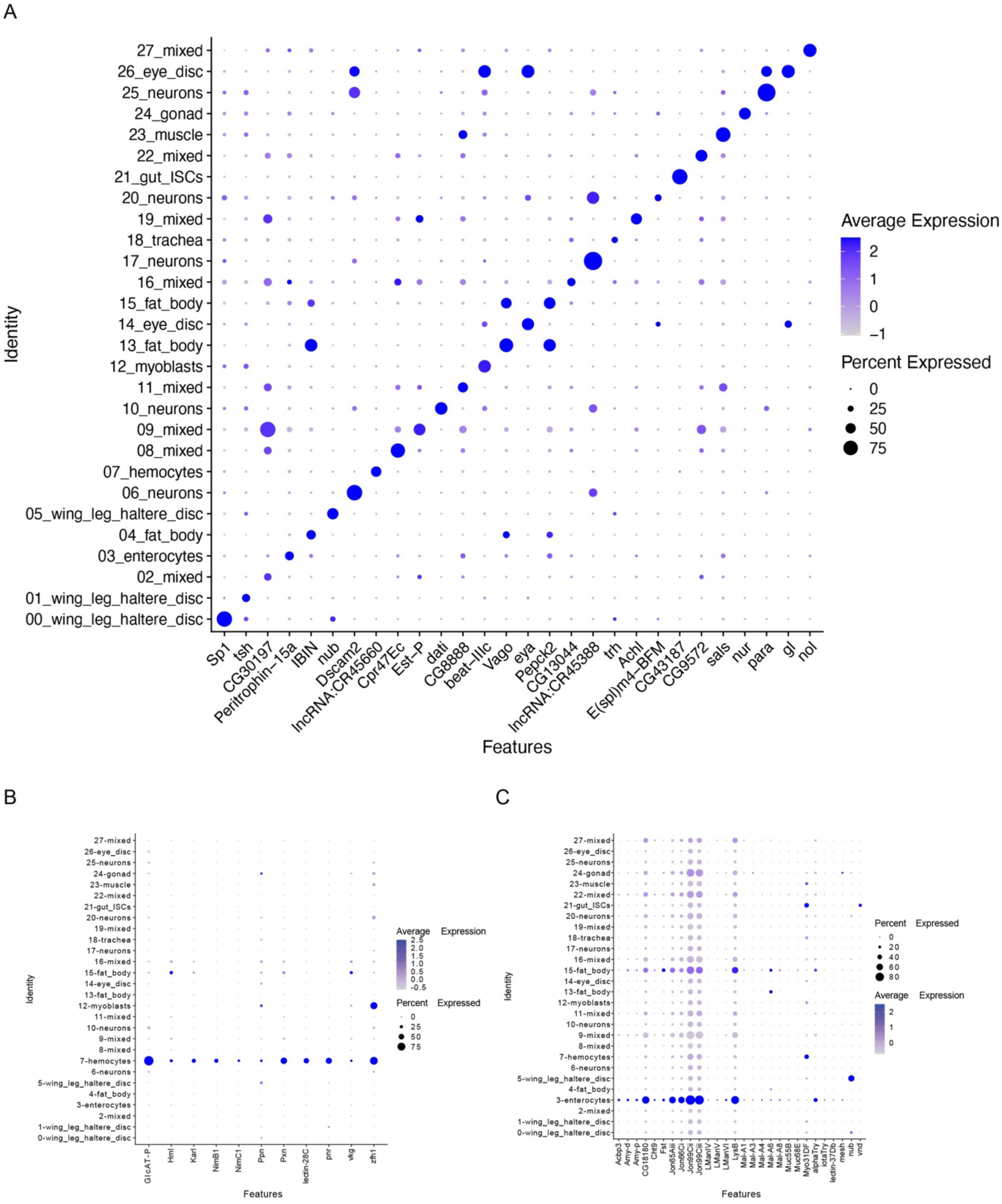
Related to Figure 4. (A-C) Dot plots showing expression levels and percentage of cells expressing the top marker gene in each cluster based on average expression (logFC). Color gradient of the dot represents the expression level, while the size represents percentage of cells expressing the marker gene per cluster. (A) Top marker gene of all clusters. (B) Hemocyte marker genes. (C) Gut enterocyte marker genes.

**Supplemental Figure 9:**
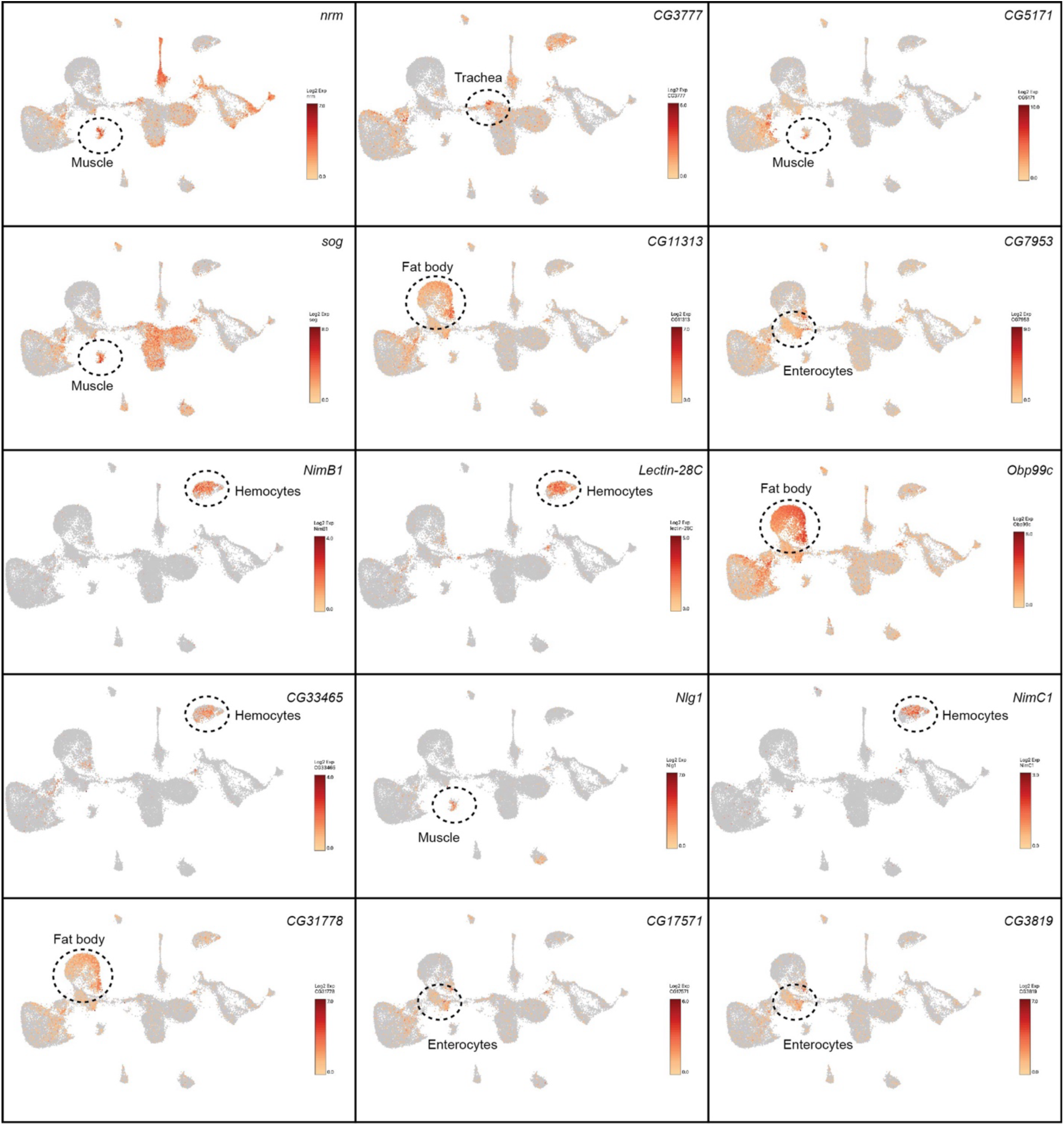
Related to Figure 4. UMAP of specific Log2 expression of selected genes encoding high-confidence tissue-specific secreted proteins.

**Supplemental Figure 10:**
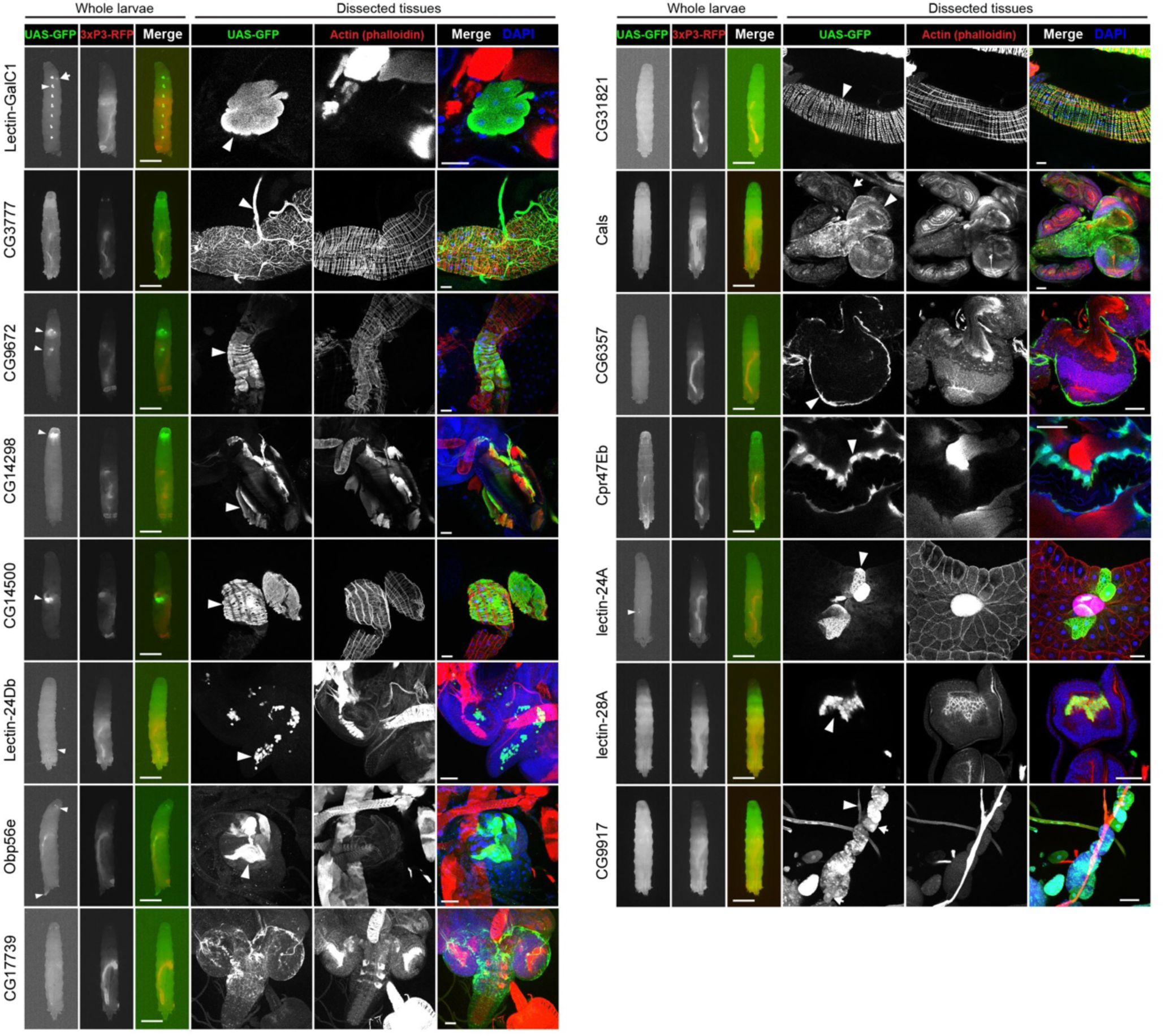
Related to Figure 5. Widefield fluorescence microscopy images of whole 3^rd^ instar larvae, or confocal microscopy of dissected tissues from 3^rd^ instar larvae, expressing *UAS-GFP* (green) under the control of a *gene-T2A-Gal4* transgene. GFP fluorescence was imaged in whole heat-killed larvae or using anti-GFP-488 (green) antibodies in dissected tissues. Also shown is *3xP3-RFP* (red) marker fluorescence for whole larvae, and phalloidin (red) and DAPI (blue) for confocal images of dissected tissues. For confocal images, the imaged tissue(s) are: Lectin-GalC1 (arrowhead indicates oenocytes), CG3777 (arrowhead indicates trachea associated with gut), CG9672 (arrowhead indicates gut enterocytes), CG14298 (arrowhead indicates mouth hooks), CG14500 (arrowhead indicates gut enterocytes), Lectin-24Db (arrowhead indicates hemocytes associated with eye disc), Obp56e (arrowhead indicates spiracles), CG17739 (brain), CG31821 (arrowhead indicates visceral muscle), Cals (arrowhead indicates brain and arrow indicates imaginal discs), CG6357 (arrowhead indicates perineural glia), Cpr47Eb (arrowhead indicates epidermis), lectin-24A (arrowhead indicates fat body cells adjacent to gonad), lectin-28A (arrowhead indicates hemocytes associated with eye disc), CG9917 (arrowhead indicates heart tube, arrows indicate pericardial nephrocytes). Scale bars are 1mm for whole larvae and 50µm for dissected tissues. All confocal images are projections except *Lectin-GalC1-T2A-Gal4*, *Cals-T2A-Gal4*, *CG6357-T2A-Gal4*, *Cpr47Eb-T2A-Gal4*, and *lectin-28A-T2A-Gal4*, which are slices.

**Supplemental Figure 11:**
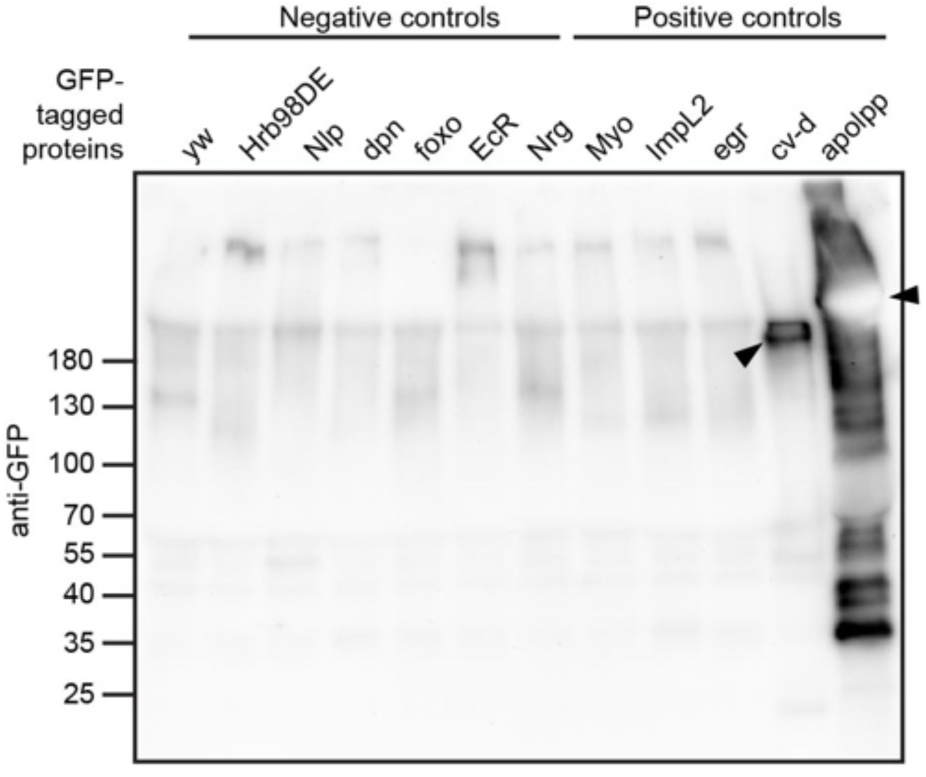
Related to Figure 5. Western blot of 3^rd^ instar larval hemolymph collected from GFP-tag lines. Negative controls are intracellular GFP-tagged proteins and positive controls are GFP-tagged known hemolymph proteins. Arrowheads indicate bands at the expected molecular weight (197.1 kDa = cv-d; 330 kDa = apoL1).

