## Supplemental File 1 for "Multi-omic mapping of *Drosophila* protein secretomes reveals tissue-specific origins and inter-organ trafficking"

GFP-TurboID-ER

MKLCILLAVVAFVGLSLGESKGEELFTGVVPILVELDGDVNGHKFSVRGEGEGDATNGKLTLKFICTTGKLPVPWPTLVTTLTYGVQCFSRYPDHMKQHDFFKSAMPEGYVQERTISFKDDGTYKTRAEVKFEGDTLVNRIELKGIDFKEDGNILGHKLEYNFNSHNVYITADKQKNGIKANFKIRHNVEDGSVQLADHYQQNTPIGDGPVLLPDNHYLSTQSVLSKDPNEKRDHMVLLEFVTAAGITLGMDELYKTGGSGGSGGSGGSGGSGGSGGKDNTVPLKLIALLANGEFHSGEQLGETLGMSRAAINKHIQTLRDWGVDVFTVPGKGYSLPEPIPLLNAKQILGQLDGGSVAVLPVVDSTNQYLLDRIGELKSGDACIAEYQQAGRGSRGRKWFSPFGANLYLSMFWRLKRGPAAIGLGPVIGIVMAEALRKLGADKVRVKWPNDLYLQDRKLAGILVELAGITGDAAQIVIGAGINVAMRRVEESVVNQGWITLQEAGINLDRNTLAATLIRELRAALELFEQEGLAPYLPRWEKLDNFINRPVKLIIGDKEIFGISRGIDKQGALLLEQDGVIKPWMGGEISLRSAEKKDEL
